# Differentiation-coupled intron retention reveals a candidate NKG2D-TR-like isoform at the murine *Klrk1* locus

**DOI:** 10.64898/2026.08.21.746301

**Authors:** Ibrahim H. Topkaya, Mobin Karimi

**Author notes:** CORRESPONDENCE: Mobin Karimi, Assistant Professor of Immunology and Microbiology, SUNY, Upstate Medical University, 766 Irving Ave., Weiskotten Hall Suite 2281, Syracuse, New York, 13210, USA.

## Abstract

NKG2D (encoded by *KLRK1* in humans and *Klrk1* in mice) is an activating receptor expressed by cytotoxic lymphocytes. In humans, NKG2D signaling is regulated post-transcriptionally: activated T cells retain intron 4 of *KLRK1* to generate NKG2D-TR, a truncated dominant-negative isoform that limits receptor signaling. Whether mice, the principal preclinical model for NKG2D-directed therapies, possess an analogous regulatory mechanism remains unknown. Across four murine RNA-seq datasets comprising 50 samples and spanning T-cell differentiation, graft-versus-host disease, and acute and chronic LCMV infection, we examined the retained-intron isoform *Klrk1-203*. The transcript retains the canonical start codon, while its predicted stop codon lies within the terminal exon downstream of the final exon-exon junction, suggesting that it may escape nonsense-mediated decay. If translated, *Klrk1-203* is predicted to encode a truncated product that retains the cytoplasmic and transmembrane domains but lacks most of the ligand-binding ectodomain. This predicted architecture resembles human NKG2D-TR, although the murine product contains a short C-terminal sequence encoded by the retained intron. *Klrk1-203* was below the detection limit in unchallenged naive and early-effector T cells but was induced in differentiated effector and memory populations, reaching approximately one-fifth of total *Klrk1* transcripts in one effector-memory sample. Read-level analyses independently demonstrated increased intron 4 retention with differentiation; however, short-read sequencing cannot fully distinguish *Klrk1-203* from the co-retained *Klrk1-204* transcript, making isoform-specific abundance dependent on model-based quantification. An independent coding-potential algorithm classified *Klrk1-203* as non-coding, providing an important counterpoint to the structural predictions. Together, these findings identify *Klrk1-203* as a candidate NMD-resistant, differentiation-associated regulator of murine NKG2D and a potential counterpart of human NKG2D-TR that warrants experimental validation.

## 1 Introduction

NKG2D (Natural Killer Group 2, member D; encoded by *KLRK1* in humans and *Klrk1* in mice) is a key activating receptor at the interface of innate and adaptive immunity (1–3). Expressed by NK cells, CD8⁺ T cells, NKT cells, and subsets of γδ T cells, NKG2D lacks an intrinsic intracellular signaling motif and instead transmits signals through the adaptor molecules DAP10 and DAP12 (1, 4, 5). Rather than recognizing foreign antigens, NKG2D detects self-encoded, MHC class I-like stress ligands—including MICA/B and ULBP1–6 in humans and RAE-1, H60, and MULT1 in mice—that are upregulated by transformed, virally infected, or damaged cells. This stress-surveillance system enables cytotoxic lymphocytes to recognize and eliminate distressed cells (1, 4).

Because NKG2D output must be strong enough to clear dangerous cells yet restrained enough to spare healthy tissue, its activity is tightly regulated as T cells progress through activation and differentiation into effector and memory states (6). This balance becomes clinically decisive after allogeneic hematopoietic stem cell transplantation, where NKG2D on donor CD8^+^ T cells drives both the beneficial graft-versus-tumor (GVT) response and the damaging graft-versus-host disease (GVHD) that limits transplant success (7, 8). The same receptor is currently being pursued as a therapeutic target in cancer and transplantation (5, 9), making the mechanisms that govern its activity a question of direct translational importance.

One pervasive but easily overlooked way immune cells tune such receptors is alternative splicing, which reshapes the transcriptome during T cell activation and allows a single gene to produce functionally distinct messages(10). Activation-induced splicing controls key signaling components in T cells, including MALT1 and the CELF2-MKK7 axis(11, 12) It can also generate isoforms that actively shape immune function, as shown for a CD28-driven PKM splice variant that supports CD8^+^ T cell metabolism(13). Intron retention is a particularly widespread form of this regulation. Instead of being removed, an intron is retained in the mature transcript, a mechanism that broadly tunes mammalian transcriptomes(14), and operates dynamically during CD4^+^ T cell activation and granulocyte differentiation(15).

Most intron-retaining transcripts carry a premature termination codon and are subsequently cleared by nonsense-mediated decay (NMD), the surveillance pathway that degrades aberrant mRNAs(16). A transcript escapes this fate only under specific structural conditions. According to the 50-nucleotide rule, if a stop codon is located in the last exon (or fewer than about 50 nucleotides upstream of the final exon-exon junction) it escapes detection as premature, sparing the transcript and allowing translation(17). When such an NMD-escaping isoform maintains an intact open reading frame, it can yield a functional protein. Human *KLRK1* is a clear example: activated human T cells retain intron 4 to generate NKG2D-TR, a truncated isoform that escapes degradation and acts as a dominant-negative brake on NKG2D signaling(18), establishing intron retention as a genuine post-transcriptional control point at the NKG2D locus.

Whether mice possess an equivalent brake remains unknown. This represents a critical knowledge gap, as the mouse is the principal preclinical platform for NKG2D-targeted GVHD and cancer immunotherapies (4, 5, 8). Murine NKG2D regulation has so far been described mainly through the NKG2D-Long and NKG2D-Short isoforms, which arise from alternative promoter use and translation (4). Yet the mouse *Klrk1* annotation already contains intron-retaining transcripts, including *Klrk1-203* (ENSMUST00000137660) (19). Its expression across naive, effector, and memory T cells, its sensitivity to NMD, its coding potential, and its relevance to GVHD and infection have not yet been characterized. Without knowing whether murine *Klrk1* carries an analogous rheostat, results from murine NKG2D-blockade studies cannot be confidently extrapolated to human therapy.

To close this gap, we quantified transcript-level RNA-seq data across four independent murine datasets spanning allogeneic GVHD and acute and chronic LCMV infection (conditions that drive robust T cell differentiation). We combined this transcriptomic approach with several supporting analyses, including a housekeeping-gene negative control, splice-junction visualization, independent coding-potential assessment, and structural protein-domain mapping. We set out to determine when the retained-intron isoform *Klrk1-203* is expressed, whether it can escape NMD to encode a truncated product, and whether it represents a candidate, NKG2D-TR-like post-transcriptional modulator of murine NKG2D.

## 2 Materials and Methods

### 2.1 Study Design

In this study we utilized a computational, isoform-level transcriptomic approach to characterize how usage of the murine *Klrk1* (NKG2D) retained-intron isoform *Klrk1-203* changes across stages of T cell differentiation and in allogeneic graft-versus-host disease (GVHD), as well as to evaluate its post-transcriptional regulation and coding potential. We performed analyses across four independent, publicly available RNA-seq datasets totaling 50 biological samples. These were supported by orthogonal validation layers, including a housekeeping gene negative control, splice junction visualization, independent coding potential assessment, nonsense-mediated decay (NMD) susceptibility prediction, *in silico* structural protein-domain mapping, and promoter transcription-factor motif analysis. The primary GVHD analysis was performed on a publicly available chronic GVHD CD4^+^ T cell RNA-seq dataset (GSE147371)

(20). To establish healthy baselines and trace isoform dynamics throughout the naive-to-effector-to-memory cascade, we additionally analyzed FACS-sorted CD4^+^ and CD8^+^ T cell populations from the Immunological Genome Project (ImmGen GSE109125) (21). Independent validation was performed on a CD8^+^ T cell dataset generated by our laboratory, comparing wild-type and TCF-7 conditional knockout (cKO) mice in an allogeneic transplant context (GSE203167) (22, 23). Finally, to extend validation to physiological, infection-driven CD8^+^ T cell differentiation, we analyzed bulk RNA-seq samples from GSE119943 (24). All analyses were computational; no human subjects or animal protocols were required.

### 2.2 Quality Control and Transcript Quantification

We performed transcript-level quantification using Salmon (v1.10.3) in mapping-based mode with selective alignment (25). Transcriptome-only Salmon indices were built from the *Mus musculus* transcriptome (GRCm39, Ensembl release 115; 116,358 target transcripts; RefSeq accession GCF_000001635.27) (26), at three k-mer sizes matched to read length: k=31 for GSE147371 (75 bp), k=21 for GSE109125 (25 bp), and k=27 for both GSE203167 (51 bp) and GSE119943 (50 bp single-end).

To optimize mapping accuracy within each dataset, these k-mer indices were explicitly matched to their respective read lengths. All indices were generated from the exact same reference transcriptome file (Mus_musculus.GRCm39.cdna.all.fa.gz), ensuring identical transcript target sets across all analyses.

Although read lengths varied across datasets, our primary metric (the proportional contribution of retained-intron isoforms to total *Klrk1* expression) is computed strictly within each sample and is therefore robust to inter-dataset variations in sequencing depth and read length. Transcriptome-only indices were utilized rather than decoy-aware (genome plus transcriptome) indices. Index equivalence was formally validated by quantifying the same GVHD sample (SRR11389222) with both the decoy-aware index (14 GB, requiring >12 GB RAM) and the transcriptome-only index (514 MB, requiring ∼1–2 GB RAM) (27). The resulting transcript-level TPM estimates demonstrated high concordance, showing a Pearson *r* = 0.992 and *R*² = 0.985 across all 116,358 transcripts, and a Spearman *ρ* = 0.961 among the 53,468 non-zero transcripts. Specifically, *Klrk1* isoform TPM differences between the indices ranged from only 2% to 11%, yielding no qualitative changes in isoform ranking or biological interpretation. The transcriptome-only index was therefore adopted for all subsequent analyses.

All Salmon quantification runs utilized the following parameters: automatic library type detection (-l A), selective alignment validation (--validateMappings), fragment-level GC bias correction (--gcBias), and sequence-specific bias correction (--seqBias) (25). Bias correction was applied across all datasets; its inclusion is critical because retained-intron quantification is highly sensitive to systematic bias. In a preliminary test on the discovery data, omitting the bias-correction flags artificially inflated the estimate of one transcript, Klrk1-202 (ENSMUST00000095412), from 0 to 52 TPM. With bias correction uniformly applied, however, Klrk1-202 is resolved as a genuine protein-coding signal rather than an artifact: its alternative first exon demonstrates read coverage and splice-junction support, and it is retained in the total-Klrk1 quantification.

Because Klrk1-202 shares its entire exon sequence with Klrk1-205 and no individual read can definitively distinguish between the two, its abundance is not estimated independently. Instead, it is reported within a combined Klrk1-205/202 unit (see Isoform reporting units).

Library chemistry differed across datasets and is relevant to retained-intron quantification. GSE109125 (Smart-seq2) and GSE203167 used poly(A)-based capture, whereas GSE147371 libraries were prepared from rRNA-depleted total RNA (KAPA HyperPrep with RiboErase). Because rRNA-depleted libraries retain nascent and incompletely processed transcripts that poly(A) selection largely removes, they are expected to carry a higher intronic background. The lower mapping rates observed for GSE147371 against a transcriptome-only index (40.5–61.8%) are consistent with this. Notably, the retained-intron fraction at Klrk1 was not higher in the rRNA-depleted dataset: GVHD CD4⁺ Tem samples reached 28.0%, compared with 34.4% in poly(A)-selected ImmGen effector memory cells. Library chemistry therefore does not account for the retained-intron signal reported here.

For the GVHD dataset (GSE147371), the two technical replicate FASTQ files (derived from separate HiSeq lanes) for each biological sample were provided simultaneously to Salmon’s –1 and –2 input arguments. This allowed Salmon to internally merge the technical replicates during the quantification step, eliminating the need for pre-processing concatenation. Mapping rates for these merged GVHD samples ranged from 40.5% to 61.8% (Table 1). ImmGen samples, which were each sequenced in a single run, achieved mapping rates between 46.0% and 77.1%, consistent with the SmartSeq2 protocol’s poly (A)-enriched library preparation. Finally, the GSE203167 samples achieved mapping rates of 78% to 86%, reflecting their longer read length and higher library quality (22).

**Table 1.** Sample structure and mapping metrics for the GSE147371 chronic GVHD dataset. Bulk RNA-seq data from FACS-purified allogeneic CD4+ T cells isolated 14 days post-transplant in a murine chronic graft-versus-host disease (GVHD) model. Naive (Tn) and effector memory (Tem) phenotypes were evaluated. Each biological sample was sequenced across two technical replicate lanes (HiSeq 4000), which were merged dynamically during Salmon quantification. Total fragments and alignment mapping rates are reported per biological replicate (n=1 Tn, n=3 Tem).

| GEO Accession | Condition | SRR Run 1 | SRR Run 2 | Fragments | Mapping Rate |
| --- | --- | --- | --- | --- | --- |
| GSM4427964 | CD4 Tem | SRR11389222 | SRR11389223 | 77.8M | 40.5% |
| GSM4427966 | CD4 Tem | SRR11389224 | SRR11389225 | 72.7M | 61.6% |
| GSM4427968 | CD4 Tem | SRR11389226 | SRR11389227 | 72.3M | 61.8% |
| GSM4427970 | CD4 Tn | SRR11389228 | SRR11389229 | 81.3M | 47.6% |

### 2.3 Isoform Proportion Analysis

Transcript-level annotations for *Klrk1* (ENSMUSG00000030149) were retrieved from Ensembl BioMart release 115 (28, 29) (Table 2). Transcript-level TPM (transcripts per million) values from Salmon were extracted, and isoform proportions were calculated as the fraction of each transcript’s TPM relative to the total gene TPM. This compositional approach captures shifts in isoform usage independent of overall gene expression changes. No minimum TPM threshold was applied prior to isoform proportion calculations; the retained-intron fraction was computed as retained-intron TPM / total *Klrk1* TPM per sample, with samples in which total *Klrk1* TPM = 0 assigned a retained-intron percentage (RI%) of 0 to avoid division by zero. Because this study focused on a single, well-characterized locus rather than a genome-wide differential expression screen, transcript-level filtering was unnecessary. Furthermore, such filtering could have introduced selection bias against low-expression naive samples, which are themselves a key biological finding of this work. Comparisons were made across all conditions spanning the four datasets (Table 5).

**Table 2.**
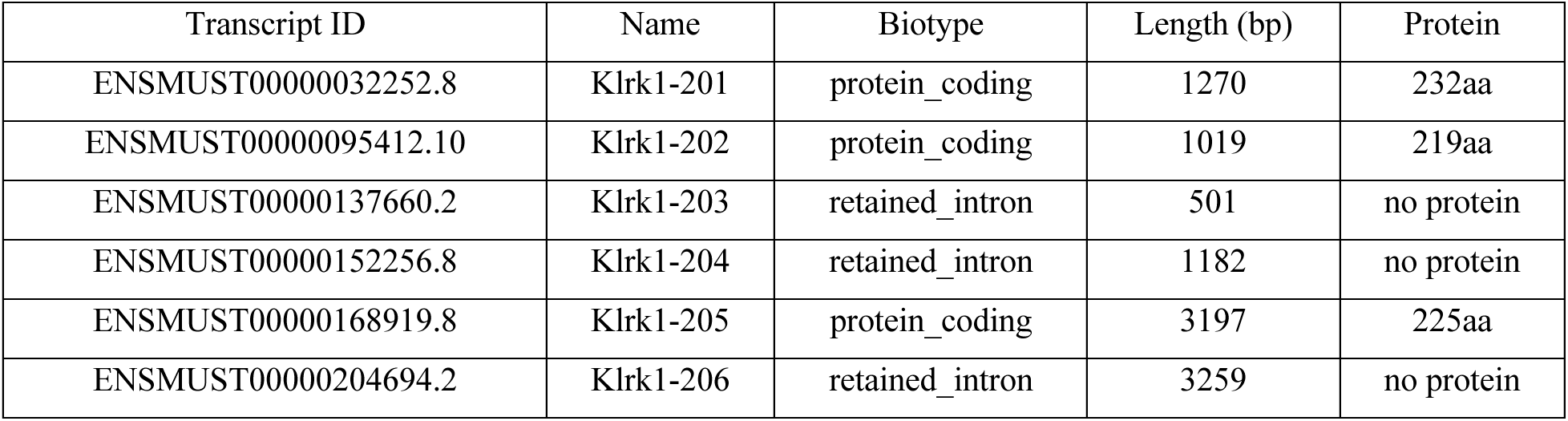
Characteristics of annotated murine *Klrk1* transcripts. Summary of the six murine *Klrk1* (NKG2D, ENSMUSG00000030149) transcripts currently annotated in Ensembl release 115 (GRCm39). Transcripts are classified by biotype (protein-coding versus retained-intron), total nucleotide length, and translated protein product length. *Klrk1-202* shares its entire exon sequence with *Klrk1-205*; because no short sequencing read can definitively separate the two, they are reported within a combined *Klrk1-205/202* unit in all subsequent quantifications. Transcript identifiers and versions are as annotated in Ensembl release 115, the release used to build the quantification index. The Klrk1 annotation has been updated in subsequent releases, including the addition of a further transcript model; all analyses reported here use the release 115 annotation throughout.

| Transcript ID | Name | Biotype | Length (bp) | Protein |
| --- | --- | --- | --- | --- |
| ENSMUST00000032252.8 | Klrk1-201 | protein_coding | 1270 | 232aa |
| ENSMUST00000095412.10 | Klrk1-202 | protein_coding | 1019 | 219aa |
| ENSMUST00000137660.2 | Klrk1-203 | retained_intron | 501 | no protein |
| ENSMUST00000152256.8 | Klrk1-204 | retained_intron | 1182 | no protein |
| ENSMUST00000168919.8 | Klrk1-205 | protein_coding | 3197 | 225aa |
| ENSMUST00000204694.2 | Klrk1-206 | retained_intron | 3259 | no protein |

### 2.4 *Klrk1* isoform reporting units

Ensembl annotates six *Klrk1* transcripts (GRCm39): three protein-coding (*Klrk1-201*, *-202*, *-205*) and three retained-intron (*Klrk1-203*, *-204*, *-206*). Because these transcripts are assembled largely from the same exons, short sequencing reads cannot always be unambiguously assigned to a single isoform. Consequently, reporting units were defined directly from the annotation itself. Mapping the locus into regions of constant transcript membership revealed that *Klrk1-204*, *-205*, and *-206* each carry a large, transcript-exclusive region (784, 2,113, and 2,865 bp, respectively) and are quantified individually. *Klrk1-203* possesses no exclusive base, but its retained intron-4 segment (chr6:129,593,531–129,593,631) is shared only with *Klrk1-204*. Because *Klrk1-204* additionally retains intron 3 (chr6:129,593,737–129,594,445, 709 bp), which *Klrk1-203* splices out, the intron-3 splice junction effectively separates the two at the read level, allowing *Klrk1-203* to be reported individually. *Klrk1-201* is distinguished by its canonical first exon. *Klrk1-202*, however, is a complete exonic subset of *Klrk1-205* (no *Klrk1-202* base lies outside *Klrk1-205*) and is linked to it by an alternative first exon and a splice junction shared only by the two.

As a result, no individual read positively distinguishes *Klrk1-202* from *Klrk1-205*; *Klrk1-202* is therefore not quantified on its own but is reported within a combined *Klrk1-205/202* unit. Quantities are thus expressed in five units (*Klrk1-201*, *Klrk1-205/202*, *Klrk1-203*, *Klrk1-204*, and *Klrk1-206*), with total *Klrk1* defined as the sum of all six transcripts, and the retained-intron fraction defined as (*Klrk1-203* + *Klrk1-204* + *Klrk1-206*) / total *Klrk1*. Because the split between co-retaining isoforms (notably *Klrk1-203* versus *Klrk1-204*) rests on model-based apportionment, isoform percentages are provided in tables, while *Klrk1-203* dynamics are described in the text as direction and fold-change.

### 2.5 Housekeeping Gene Negative Control

To exclude pipeline artifacts and genomic DNA contamination as explanations for the observed Klrk1 retained-intron signal, an identical isoform proportion analysis was performed on two housekeeping genes, Actb (ENSMUSG00000029580) and Gapdh (ENSMUSG00000057666), across the 31 samples of the discovery and primary-validation datasets (and, separately, across the 19 GSE119943 samples). Transcript-level TPM values for all annotated isoforms of these genes were extracted from the Salmon quant. sf files, and the proportion of total gene TPM contributed by retained-intron isoforms was calculated using the same methodology applied to Klrk1. Because Actb and Gapdh are highly expressed, constitutively spliced housekeeping genes, any substantial retained-intron signal at these loci would indicate a pipeline-level artifact rather than gene-specific intron retention (Fig. 1).

**Figure 1.**
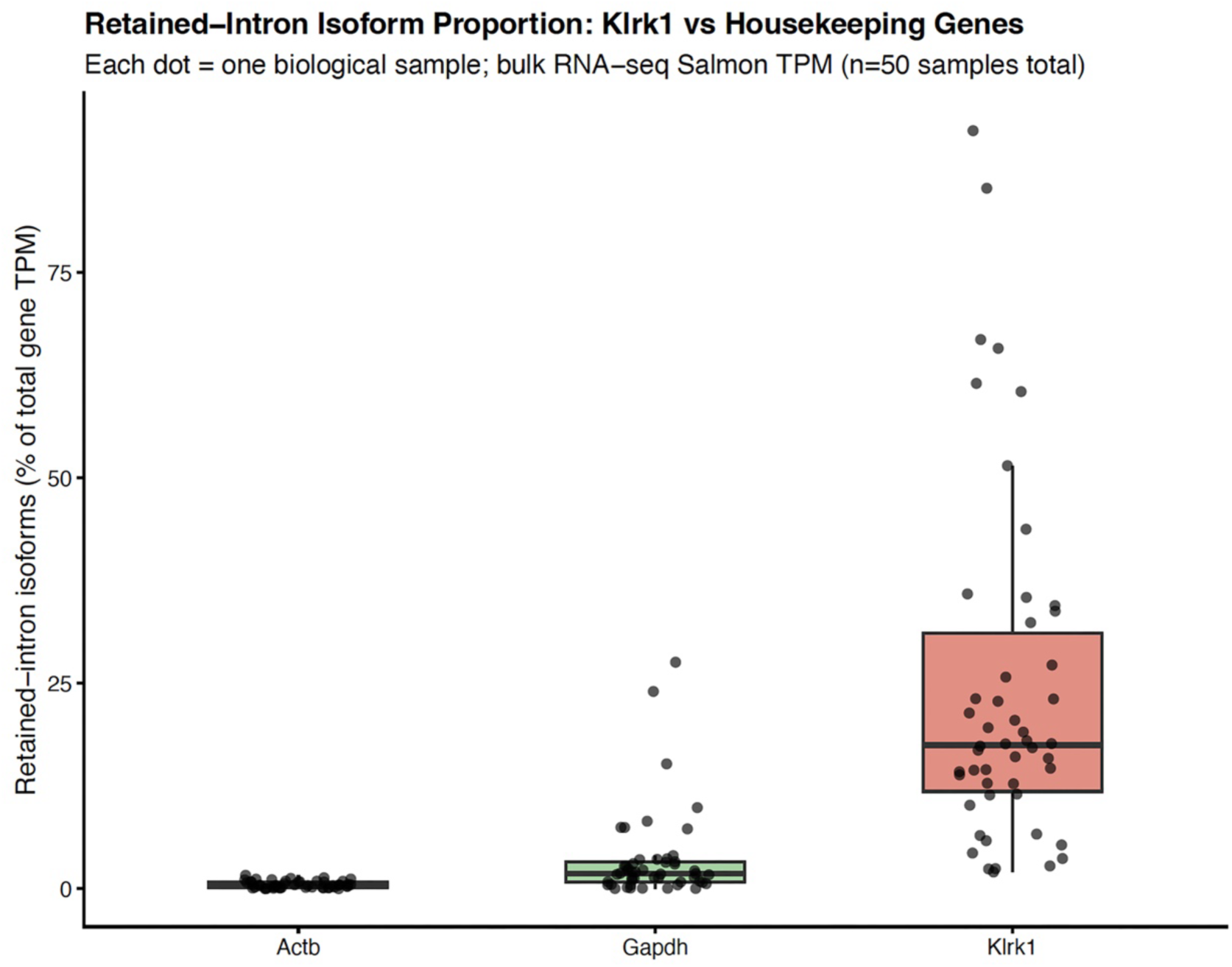
High-specificity intron retention at the *Klrk1* locus compared to housekeeping controls. The retained-intron proportion (RI%) (calculated as the sum of all annotated retained-intron transcripts divided by total gene TPM) is compared between *Klrk1* and the constitutively spliced housekeeping genes *Actb* and *Gapdh*. Data aggregate all 50 biological samples across the four independent datasets (GSE147371, GSE109125, GSE203167, GSE119943). The median RI% at *Klrk1* (17%) substantially exceeds background transcriptional/splicing noise at *Actb* (0.40%) and *Gapdh* (1.87%), confirming that *Klrk1* intron retention is a targeted biological mechanism rather than an artifact of incomplete library processing or genomic DNA contamination

### 2.6 Datasets

#### 2.6.1 Dataset 1: GSE147371 (GVHD CD4^+^ T cells, primary discovery)

Bulk RNA-seq data were obtained from the Gene Expression Omnibus (GEO) under accession GSE147371 (BioProject PRJNA614021), originally generated by Assmann *et al.* to study the glycolytic metabolism of pathogenic T cells in GVHD (20). The B10.D2 donor strain used in the GVHD model (Jackson Laboratory stock #000463) is a congenic strain in which the DBA/2-derived H2 complex was introgressed onto a C57BL/10Sn background. C57BL/10 and C57BL/6 are related but distinct inbred strains, so the ImmGen C57BL/6J profiles provide an approximate rather than strain-matched resting reference for the GVHD donor compartment (30). The lethally irradiated BALB/c (H-2d, Jackson Laboratory stock #555) recipients were conditioned with 850 cGy total body irradiation, delivered in two doses three hours apart on day −1, and then reconstituted with 15 million unfractionated splenocytes and 8 million bone marrow cells injected via tail vein on day 0. FACS-purified allogeneic CD4^+^ T cells were isolated from spleens at day 14 post-transplant and sorted into naive (Tn; CD44^lo^ CCR7^+^) and effector memory (Tem; CD44^hi^ CCR7^-^) populations (20). To ensure sufficient RNA yield (>0.5 million cells per sample), cells of each phenotype were pooled from multiple study animals. RNA was extracted from these pooled samples, yielding one biological replicate for Tn (n=1) and three biological replicates for Tem (n=3). Sequencing was performed on the Illumina HiSeq 4000 platform, generating 75 bp paired-end reads (Table 1).

#### 2.6.2 Dataset 2: GSE109125 (ImmGen, healthy controls and CD8^+^ differentiation series)

To establish resting baselines and characterize differentiation-associated isoform usage, we utilized Ultra-Low-Input RNA-seq data from the ImmGen compendium (GEO accession GSE109125). Naive populations were taken from unchallenged mice, whereas the effector and memory populations in this compendium were generated in an LCMV infection model; we therefore treat the ImmGen comparison as a naive-versus-antigen-experienced contrast rather than a steady-state series, and note that it shares an infection-driven differentiation axis with GSE119943 rather than being fully orthogonal to it (21). Naive CD4^+^ T cells (n=4 biological replicates; sorting: CD4^+^ CD8^−^ TCRβ^hi^ CD62L^hi^ CD44^lo^ CD25^−^ Dump–)(31) and naive CD8^+^ T cells (n=2 biological replicates; sorting: CD4^−^ CD8^+^ TCRβ^hi^ CD62L^hi^ CD44^lo^ Dump–) were selected from FACS-sorted splenic populations of healthy C57BL/6J mice.

To trace the naive-to-effector-to-memory cascade, we additionally selected CD8⁺ terminal effector cells (T8.TE, LCMV day 7; n=2), memory precursor cells (T8.MP, LCMV day 7; n=2), central memory cells (T8. Tcm, LCMV day 180; n=2), and effector memory cells (T8. Tem, LCMV day 180; n=1), together with naive P14 TCR-transgenic CD8⁺ T cells (T8.TN.P14; n=2) as an additional resting reference. Population identities and sorting strategies were taken directly from the ImmGen sample annotations in GEO.

ImmGen samples were generated using the Smart-seq2 ultra-low-input protocol (∼500– 1,000 cells per replicate) and sequenced on the Illumina NextSeq 500 platform, producing 25 bp paired-end reads. The shorter read length of the ImmGen dataset (25 bp, compared to 75 bp for GSE147371) necessitated the construction of a separate Salmon index with a reduced k-mer size (k=21 vs. k=31). The CD4+ naive sorting strategy utilized for the ImmGen dataset (CD4^+^ CD8^−^ TCRβ^hi^ CD62L^hi^ CD44^lo^ CD25^−^ Dump^−^) is closely comparable to, though not identical with, the Tn definition used in the GVHD dataset, which relied on CCR7 rather than CD62L (Table 3).

**Table 3.** Sample structure and mapping metrics for the ImmGen dataset (GSE109125). Ultra-low-input Smart-seq2 RNA-seq profiles of FACS-sorted splenic T cell populations from C57BL/6J mice. Naive CD4⁺ and CD8⁺ populations were obtained from unchallenged animals, whereas the antigen-experienced CD8⁺ populations were generated in an LCMV infection model and comprise CD8+ terminal effector cells (T8.TE, day 7), CD8+ memory precursor cells (T8.MP, day 7), CD8+ central memory cells (T8.Tcm, day 180) and CD8+ effector memory cells (T8.Tem, day 180). Naive P14 TCR-transgenic CD8⁺ cells (T8.TN.P14) serve as an additional resting reference, and naive CD4⁺ cells provide a cross-lineage baseline. Population names and sorting strategies are reported as annotated by ImmGen in GEO; the two T.4.Nve.Fem samples derive from female mice. For the LCMV-derived populations, the difference in annotated age relative to the naive samples reflects the interval since infection rather than an independent age variable. All samples were quantified using a k=21 Salmon index matched to the 25 bp read length.

| Sample | GEO Accession | SRR | Cell Type | Age/Condition | Fragment<br>s | Mapping<br>rate |
| --- | --- | --- | --- | --- | --- | --- |
| T.4.Nve.Fem.S<br>p#1 | GSM2932601 | SRR6467017 | Naïve female CD4+ | 6 wk | 13.7M | 64.8% |
| T.4.Nve.Fem.S<br>p#2 | GSM2932602 | SRR6467018 | Naïve female CD4+ | 6 wk | 11.1M | 52.6% |
| T.4.Nve.Sp#1 | GSM2932603 | SRR6467019 | CD4+ Naïve | 6 wk | 13.6M | 64.1% |
| T.4.Nve.Sp#2 | GSM2932604 | SRR6467020 | CD4+ Naïve | 6 wk | 7.6M | 59.7% |
| T.8.Nve.Sp#1 | GSM2932609 | SRR6467025 | CD8+ Naïve | 6 wk | 13.1M | 63.8% |
| T.8.Nve.Sp#2 | GSM2932610 | SRR6467026 | CD8+ Naïve | 6 wk | 8.7M | 60.7% |
| T8.TE.LCMV.d<br>7.Sp#1 | GSM2932625 | SRR6467041 | CD8+ Terminal<br>Effector | 7 wk/<br>LCMV<br>Day7 | 12.2M | 75.5% |
| T8.TE.LCMV.d<br>7.Sp#2 | GSM2932626 | SRR6467042 | CD8+ Terminal<br>Effector | 7 wk/<br>LCMV<br>Day7 | 8.3M | 73.2% |
| T8.Tem.LCMV<br>.d180.Sp#2 | GSM2932627 | SRR6467043 | CD8+ Effector<br>Memory | 32 wk/<br>LCMV<br>Day180 | 10M | 54.8% |
| T8.TN.P14.Sp#<br>1 | GSM2932628 | SRR6467044 | Naïve CD8+ P14 | 6 wk | 8.1M | 64.3% |
| T8.TN.P14.Sp#<br>2 | GSM2932629 | SRR6467045 | Naïve CD8+ P14 | 6 wk | 5.6M | 55% |
| T8.Tcm.LCMV<br>.d180.Sp#1 | GSM2932623 | SRR6467039 | CD8+ Central<br>Memory | 32 wk/<br>LCMV<br>Day 180 | 14.1M | 63.5% |
| T8.Tcm.LCMV<br>.d180.Sp#2 | GSM2932624 | SRR6467040 | CD8+ Central<br>Memory | 32 wk/<br>LCMV<br>Day 180 | 9.1M | 61.9% |
| T8.MP.LCMV.<br>d7.Sp#1 | GSM2932621 | SRR6467037 | CD8+ Memory<br>Precursor | 7 wk/<br>LCMV<br>Day 7 | 11.2M | 74.8% |
| T8.MP.LCMV.<br>d7.Sp#2 | GSM2932622 | SRR6467038 | CD8+ Memory<br>Precursor | 7 wk/<br>LCMV<br>Day 7 | 10.6M | 77.1% |

#### 2.6.3 Dataset 3: GSE203167 (Allogeneic transplant CD8^+^ T cell validation)

To independently validate the principal findings, we analyzed a CD8^+^ T cell RNA-seq dataset generated by our laboratory (GEO accession GSE203167; BioProject PRJNA837624). This dataset profiles wild-type (WT) and TCF-7 conditional knockout (cKO) CD8^+^ T cells before transplantation (Pre-Tx) and at day 7 post-allogeneic transplantation (Post-Tx D7) in a murine model. Twelve biological samples were analyzed in total: WT Pre-Tx (n=3), WT Post-Tx D7 (n=3), TCF-7 cKO Pre-Tx (n=3), and TCF-7 cKO Post-Tx D7 (n=3) (22). Sequencing was performed on the Illumina NovaSeq platform, generating 51 bp paired-end reads. The longer read length and larger sample size relative to the primary discovery dataset (GSE147371) enabled both robust transcript quantification and high-resolution splice junction visualization (see Sashimi Plot Analysis below).

#### 2.6.4 Dataset 4: GSE119943 (LCMV CD8^+^ T cell differentiation and exhaustion validation)

To extend validation to physiological, infection-driven CD8^+^ T cell differentiation, we analyzed bulk RNA-seq samples from GSE119943 (24), a study investigating the TOX-dependent regulation of CD8^+^ T cell persistence in chronic infection. The series profiles P14 TCR-transgenic CD8^+^ T cells responding to acute (LCMV-Armstrong) and chronic (LCMV-Clone 13) infection. Nineteen bulk RNA-seq samples were analyzed in total: acute LCMV-Armstrong early effector cells (day 4.5; n=3), day-7 short-lived effector cells (SLEC; KLRG1^hi^ CD127^lo^; n=3), and day-7 memory precursor effector cells (MPEC; KLRG1lo CD127^hi^; n=3); chronic LCMV-Clone 13 day-7 progenitor-like (n=2) and terminally exhausted (n=2) cells, as annotated in GEO; and empty-vector (pMIG) control progenitor-like (n=3) and terminally exhausted (n=3) cells. Only bulk RNA-seq samples were utilized; the single-cell (10x Genomics) and ChIP-seq components of this GEO series were deliberately excluded, as 3’-biased single-cell chemistry cannot reliably quantify internal intron-4 retention. Reads were 50 bp single-end; Salmon quantification utilized a read-length-matched k=27 index in single-end mode (-r), with the same –-gcBias and –-seqBias bias correction parameters applied to all other datasets (25).

Only bulk RNA-seq samples were analysed; the single-cell (10x Genomics) and ChIP-seq components of this GEO series were excluded, as 3’-biased single-cell chemistry cannot reliably quantify internal intron-4 retention. Within the bulk RNA-seq component we selected the day-4.5 acute-infection samples together with the day-7 sorted effector and memory-precursor subsets from acute infection and the day-7 sorted progenitor-like and terminally exhausted subsets from chronic infection, so that each condition corresponds to a defined differentiation or exhaustion state. The unsorted day-7 bulk populations were not included, because they represent mixtures of these subsets, and the TOX-overexpression samples were excluded as a genetic perturbation; their empty-vector (pMIG) counterparts were retained and are reported separately.

This dataset provides a within-experiment differentiation gradient (early effector to terminal effector and memory precursor) and an additional exhaustion axis. Because the ImmGen effector and memory populations are also LCMV-derived, GSE119943 is best regarded as a replication of the same differentiation axis under controlled, within-experiment conditions rather than as a fully independent one; the GVHD and allogeneic transplant datasets provide the distinct pathological contexts.

### 2.7 Splice Junction Visualization (Sashimi Plot)

To provide direct visual evidence of splice junction usage at the *Klrk1* locus and to rule out genomic DNA contamination at the alignment level, Sashimi plots were generated for representative samples from (i) GSE203167 (one WT Pre-Tx and one WT Post-Tx D7; Figure 2) and (ii) GSE119943 (one Arm D7 SLEC and one Cl13 D7 Term-Exh; Figure 3). Genome-aligned BAM files were produced using HISAT2 (v2.2.2) against a chromosome 6-restricted GRCm39 index (built with the –-ss and –-exon options, utilizing the Ensembl 115 GTF annotation to provide known splice-site information). HISAT2 was selected after STAR (v2.7.11b) was found to be non-functional on the macOS arm64 platform used for analysis. The resulting BAM files were sorted and indexed with SAMtools (v1.22.1), and Sashimi plots restricted to the *Klrk1* locus (chr6:129,583,000–129,605,000) were generated using ggsashimi (v1.1.5) alongside the Ensembl 115 GTF annotation.

**Figure 2.**
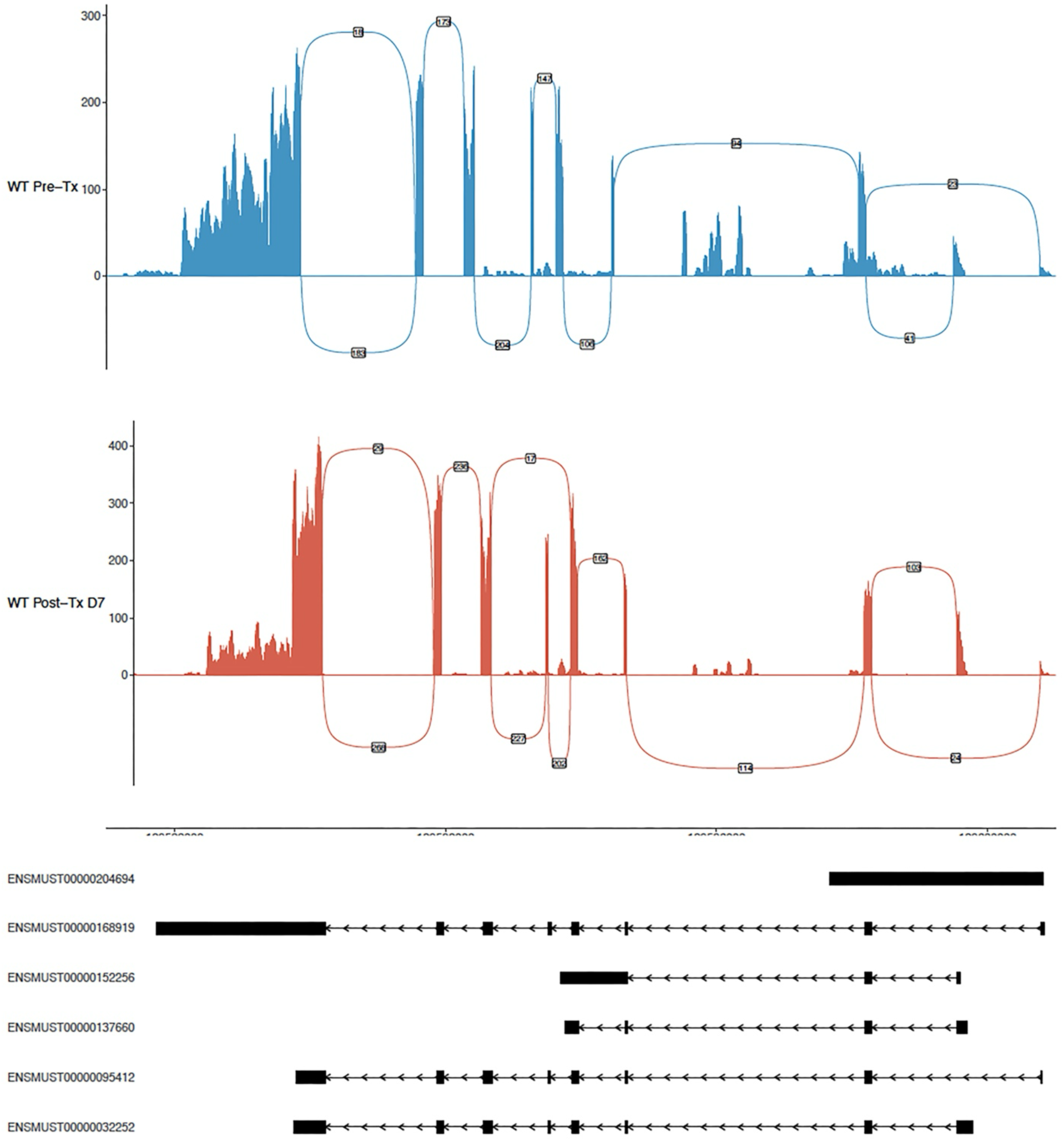
Alignment-level Sashimi plot confirmation of *Klrk1* splice-junction usage. Sashimi visualization of HISAT2-aligned reads mapped to the murine *Klrk1* locus for two representative GSE203167 samples: wild-type pre-transplant (WT Pre-Tx) and day 7 post-transplant (WT Post-Tx D7). Arcs indicate splice-junction spanning reads, while histograms display raw coverage depth. The canonical exon 4–exon 5 splicing event (which physically removes intron 4) is strongly supported by 147 reads (Pre) and 202 reads (Post-D7). Notably, direct read coverage is sustained across the retained intron 4 region, corroborating the presence of the *Klrk1-203* transcript at the raw alignment level.

**Figure 3.**
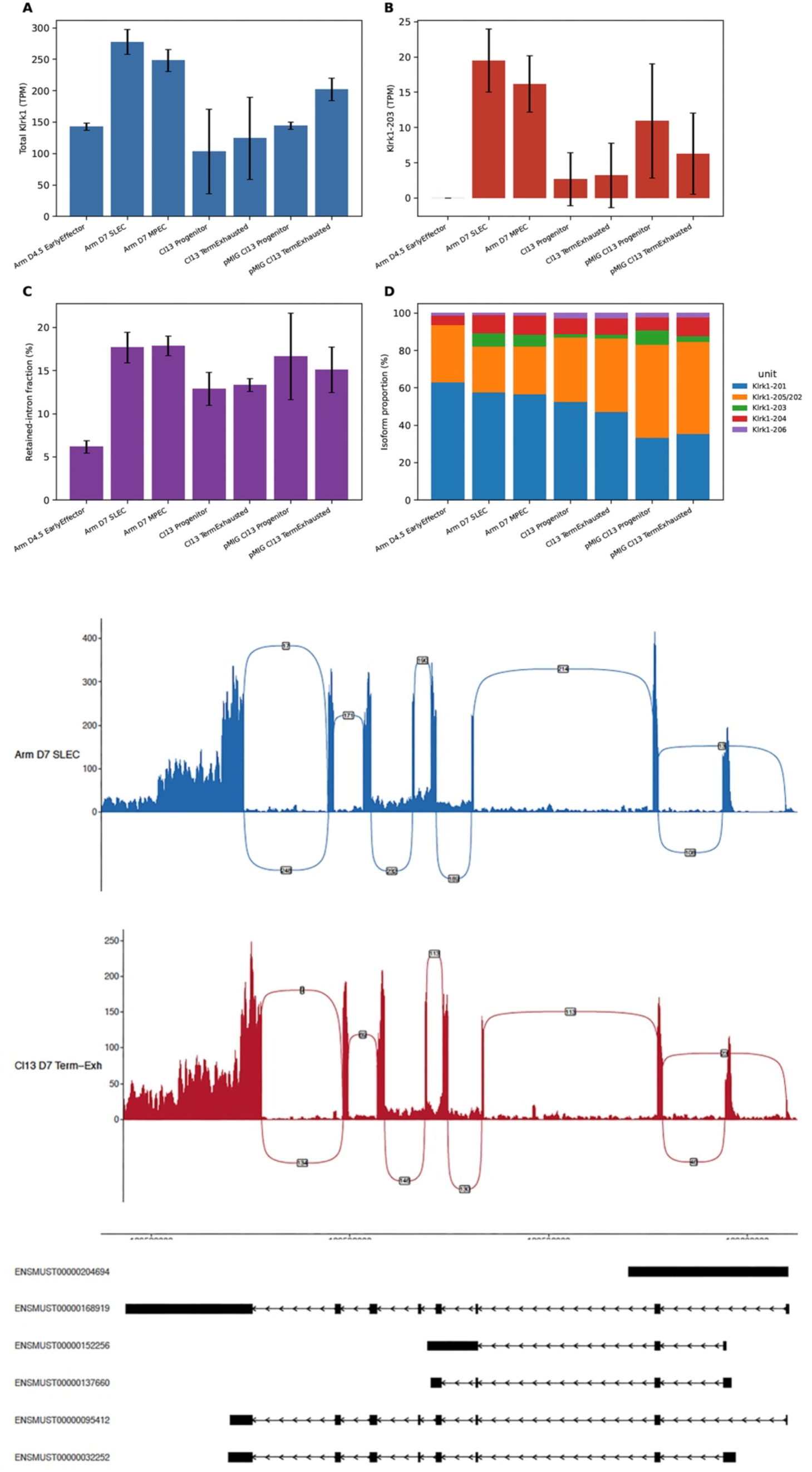
*Klrk1* splicing dynamics during acute viral differentiation and chronic exhaustion in GSE119943 dataset. Transcriptomic evaluation of P14 CD8+ T cells across acute (LCMV-Armstrong) and chronic (LCMV-Clone 13) infection trajectories (GSE119943). **(A)** Total *Klrk1* TPM expression (mean ± SD). **(B)** Absolute *Klrk1-203* expression (mean ± SD), showing a strict absence at day 4.5 and subsequent induction. **(C)** Total retained-intron fraction (% of total *Klrk1*). **(D)** *Klrk1* isoform proportions broken down by the five reporting units. Note the prominent induction of *Klrk1-203* tightly coupled to day 7 SLEC and MPEC maturation. **(E)** Sashimi plot of representative acute effector (Arm day-7 SLEC) and chronically exhausted (Cl13 day-7 terminally exhausted) samples, visually confirming locus-wide splice-junction usage and intronic coverage (5’ to 3’, right to left).

### 2.8 Junction Read Counting and Percent Intron Retention (PIR) Analysis

To independently quantify intron retention beyond Sashimi plot visualization, splice junction reads were extracted from HISAT2-aligned BAM files using custom Python scripts in conjunction with SAMtools (v1.22.1). For each *Klrk1* splice junction, reads containing an N CIGAR operation that spanned the annotated intron boundaries (with a ±10 bp tolerance) were counted. Mean coverage across each *Klrk1* exon and across the retained intron 4 region was computed using samtools depth (Table 4). Percent Intron Retention (PIR) at intron 4 was calculated as *PIR* = *retention_reads* / (*retention_reads* + *E4|E5_junction_reads*), where *retention_reads* were reads contiguously spanning the exon 4–intron 4 and intron 4–exon 5 boundaries, and *E4|E5_junction_reads* were reads with an N (skip) CIGAR operation spanning intron 4. PIR was computed for the two representative GSE203167 samples (one WT Pre-Tx, one WT Post-Tx D7); the transcript-level Salmon *Klrk1-203* fraction provided an independent, concordant estimate in these same samples. GSE147371 and GSE109125 reads were aligned to a chromosome 6-restricted GRCm39 HISAT2 index (the same index utilized for the Sashimi analysis). Intron 4 retention was quantified from reads spanning the exon 4/intron 4 and intron 4/exon 5 boundaries relative to exon 4–exon 5 spliced reads (PIR). Because the 25 bp ImmGen reads cannot be reliably scored as junction-spanning, the ImmGen dataset utilized a read-length-independent intron 4 coverage ratio (*Klrk1-203*-retained proximal intron versus constitutive exon 4). The junction metric strictly measures overall intron 4 retention; isoform-specific (*Klrk1-203*) attribution is derived from Salmon and the *Klrk1-203*-versus-*Klrk1-204/206* decomposition, as both *Klrk1-203* and *Klrk1-204* cross the exon 4/intron 4 boundary indistinguishably.

**Table 4.** Read depth support across diagnostic *Klrk1* genomic regions. Mean sequencing read depth (calculated via samtools depth) across diagnostic, transcript-specific genomic coordinates within the *Klrk1* locus. Data are shown for two representative, bias-corrected samples (WT Post-Tx D7 from GSE203167, and CD4^+^ Tem_1 from GSE147371). Mean depth over the intron-4 segment (shared by Klrk1-203 and Klrk1-204) exceeds that over the Klrk1-204-exclusive intron-3 segment in both samples (1.9-fold and 3.7-fold), which is consistent with a contribution from Klrk1-203 beyond Klrk1-204 alone. Because the two regions differ substantially in length and only two samples are shown, this comparison is indicative rather than quantitative.

| Diagnostic region | Transcript(s) | Region type | chr6 coords | mean depth: WT_Post7 (GSE203167) | mean depth: Tem_1 (GSE147371) |
| --- | --- | --- | --- | --- | --- |
| Canonical first exon | 201/203/206 | shared | 129599545-129599735 | 19.9 | 24.6 |
| Alternative first exon | 202/205/206 | shared | 129600774-129600804 | 12.9 | 6.4 |
| 205-exclusive 3' tail | 205 | EXCLUSIVE | 129587286-129589376 | 26.3 | 28.6 |
| 206-exclusive exon | 206 | EXCLUSIVE | 129599736-129600773 | 0.1 | 1.2 |
| Intron-3 retention | 204 | EXCLUSIVE | 129593737-129594445 | 1.5 | 10.1 |
| Intron-4 retention segment | 203/204 | shared | 129593531-129593631 | 5.6 | 19.7 |

### 2.9 Independent Coding Potential Assessment

To independently evaluate the non-coding annotation of *Klrk1-203* (ENSMUST00000137660), the Coding Potential Assessment Tool (CPAT, v3.0.5) was applied to the cDNA sequence retrieved from the exact same reference transcriptome FASTA file used for Salmon quantification. The mouse-specific logit model and hexamer frequency table (Mouse_logitModel. RData and Mouse_Hexamer.tsv) provided by the CPAT developers were utilized. Transcripts scoring below the predefined, mouse-specific cutoff of 0.44 were classified as non-coding.

### 2.10 NMD Susceptibility Prediction

NMD susceptibility was assessed under two complementary translation initiation scenarios. First, we mapped the canonical *Klrk1-201* start codon (genomic position chr6:129,599,534–129,599,536, GRCm39, minus strand) to the transcript coordinates of ENSMUST00000137660. Because *Klrk1-203* shares its upstream exon architecture with *Klrk1-201*, this ATG was anticipated to be present within the *Klrk1-203* transcript. Translation was modeled in-frame from this canonical ATG to identify the first downstream stop codon. Second, an unbiased three-frame open reading frame (ORF) scan was performed on the full 501-nt *Klrk1-203* sequence to identify the longest ORF independently of any prior annotation. The Kozak context surrounding each candidate start codon was evaluated using the consensus criteria of a purine at position −3 and a guanine at position +4. For each candidate ORF, the position of the in-frame stop codon was compared to the last exon-exon junction (EEJ3, transcript position 296), and the canonical 50-nucleotide rule was applied: stop codons lying more than 50 nucleotides upstream of the last EEJ classify the transcript as NMD-susceptible, while stop codons located in the last exon equivalent or within 50 nt of the last EEJ predict NMD resistance. All NMD predictions are *in silico* and do not substitute for experimental demonstration of transcript stability or translation.

### 2.11 *In Silico* Structural Mapping of the Predicted Truncated Product

ENSMUST00000137660 (*Klrk1-203*) is a 501-nucleotide, four-exon transcript. Its first three exons (transcript positions 1–148, 149–256, and 257–295) are shared with the canonical *Klrk1-201*. Its fourth, terminal exon (positions 296–501) extends from canonical exon 4 directly into intron 4, retaining the first ∼101 nucleotides of that intron before the transcript ends. Because it terminates within the retained intron 4 rather than continuing through the canonical downstream exons, *Klrk1-203* (501nt) is shorter than canonical *Klrk1-201* (1,270nt) despite retaining intronic sequence. Structurally, this retained intron lies exactly at the boundary between the transmembrane and ligand-binding domains. The position of the predicted stop codon relative to the final exon-exon junction was examined to assess NMD susceptibility: because *Klrk1-203* terminates within its terminal (fourth) exon, downstream of the last exon-exon junction, its stop codon is not expected to be recognized as premature under the 50-nucleotide rule. Consequently, the transcript is predicted to escape nonsense-mediated decay despite retaining intronic sequence.

To assess the potential functional architecture of Klrk1-203 if translated, cDNA sequences for all six annotated Klrk1 transcripts were retrieved from Ensembl (GRCm39, release 115). The retained intronic sequence in ENSMUST00000137660 was mapped to the protein domain architecture of the canonical NKG2D protein (ENSMUSP00000032252, UniProtKB O54709) by aligning exon boundaries to known domain annotations: cytoplasmic region, amino acids 1–66; transmembrane helix, amino acids 67–89; extracellular region, amino acids 90–232; C-type lectin domain, amino acids 122–228. Three-frame translation of the retained intronic sequence was performed to identify premature termination codons and predict the architecture of any putative truncated product. The predicted domain architecture was compared to the previously characterized human NKG2D-TR dominant-negative isoform (8, 18). All structural inferences are computational predictions and require experimental validation.

### 2.12 *In Silico* Promoter and Transcription-Factor Motif Analysis

*In silico* promoter analysis was performed to assess whether the *Klrk1* retained-intron isoform *Klrk1-203* (ENSMUST00000137660) could be transcriptionally regulated by activation-induced transcription factors. Because *Klrk1-203* initiates on the minus strand only 88 bp downstream of the canonical *Klrk1-201* transcription start site (TSS) (ENSMUST00000032252; chr6:129,599,735, GRCm39/(29) 115), the two transcripts share a common promoter. A 2.3-kb window spanning −2000 to +288 bp relative to the canonical TSS (chr6:129,599,447– 129,601,735) was therefore retrieved using samtools faidx. Position weight matrices for NFAT family members (*Nfatc1*, MA0624.3; *Nfatc2*, MA0152.3; *NFATC3*, MA0625.3; *NFATC4*, MA1525.3; *NFAT5*, MA0606.3) and AP-1 family members (*FOS: JUN*, MA0099.3; *BATF*, MA1634.2; *BATF: JUN*, MA0462.2) were obtained from the JASPAR 2024 CORE (vertebrates) database. These were scored on both strands using Biopython log-odds position-specific scoring matrices (PSSMs; Bio.motifs) with a pseudocount of 0.5 and the local promoter nucleotide composition as background. Match significance was assigned from the score distribution at a false-positive rate of 1 × 10⁻⁴ for NFAT and 1 × 10⁻³ for AP-1 (equivalent to the FIMO *p*-value approach). Finally, NFAT: AP-1 composite elements were defined by an edge-to-edge spacing of ≤20 bp.

### 2.13 Statistical Considerations

Formal differential transcript usage (DTU) testing was applied only where replication permitted it. In the GVHD dataset (GSE147371), which includes one Tn and three Tem biological samples, DTU testing was not performed, because these frameworks require at least three biological replicates per condition to estimate within-group variance; all reported isoform proportion changes between GVHD Tn and Tem are therefore descriptive. The ImmGen dataset (GSE109125) is likewise cross-sectional, with one to four samples per population, and is reported descriptively; it provides replicated resting baselines (four naive CD4⁺, two naive CD8⁺, and two naive P14 CD8⁺ samples) against which antigen-experienced populations are compared.

DTU testing was performed on GSE203167, which provides three biological replicates in each of four conditions and therefore carries the primary inferential weight of the study. Isoform composition was compared between conditions using a Dirichlet-multinomial model of isoform-level counts across the five reporting units.

Comparisons across datasets are descriptive given the differing experimental contexts (unchallenged versus infection-driven versus post-transplant; CD4⁺ versus CD8⁺; chronic GVHD versus acute allogeneic transplant) and sequencing platforms. All computational analyses were executed on macOS (Apple M4) within a conda environment using a native ARM64 Salmon build. R (v4.5.2) with the tximeta and ggplot2 packages was used for data aggregation and visualization.

## 3 Results

### 3.1 The *Klrk1* isoform landscape

Standard gene-level quantification of *Klrk1* (which encodes the activating receptor NKG2D) obscures critical post-transcriptional regulatory mechanisms (such as intron retention) making precise, isoform-level resolution essential for understanding its dynamic control during T cell activation (32, 33). The murine *Klrk1* locus encodes six annotated transcripts that overlap extensively: three protein-coding (*Klrk1-201*, *-202*, *-205*) and three retained-intron (*Klrk1-203*, *-204*, *-206*) variants, all assembled largely from the same exons (34). This structural overlap means some transcripts cannot be unambiguously distinguished by short sequencing reads. For instance, *Klrk1-202* is a complete exonic subset of *Klrk1-205* and is therefore reported together with it as a combined *Klrk1-205/202* unit, whereas *Klrk1-204*, *-205*, and *-206* each carry large isoform-exclusive regions and can be measured directly. *Klrk1-203*, the retained-intron isoform analogous to human *NKG2D-TR* and the primary focus of this work, retains the proximal segment of intron 4 while completely splicing out intron 3.

This specific splice combination distinguishes it from *Klrk1-204* at the read level (Methods). We therefore describe *Klrk1* isoform usage across five reporting units (*Klrk1-201*, *Klrk1-205/202*, *Klrk1-203*, *Klrk1-204*, and *Klrk1-206*), express every proportion as a fraction of the total *Klrk1* pool, and define the overall retained-intron fraction as the sum of *Klrk1-203*, *-204*, and *-206*. Throughout this analysis, the most robust metric is the direction and magnitude of *Klrk1-203* induction; the exact quantitative split between *Klrk1-203* and *-204* is inherently model-dependent and is treated as such.

To characterize *Klrk1* isoform dynamics across T cell states, we analyzed four independent murine RNA-seq datasets totaling 50 biological samples (Tables 1, 2). The primary discovery dataset (GSE147371) profiled CD4^+^ naive (Tn) and effector memory (Tem) T cells from a chronic GVHD model. The Immunological Genome Project dataset (GSE109125) provided FACS-sorted CD4^+^ and CD8^+^ T cell populations spanning the full naive-to-effector-to-memory cascade in healthy C57BL/6J mice. Independent validation was performed on a CD8^+^ T cell dataset (GSE203167) generated by our laboratory, which compared wild-type and *TCF-7* conditional knockout (cKO) mice before and after allogeneic transplantation, while a fourth dataset (GSE119943) extended this validation to an orthogonal axis of infection-driven exhaustion. Each dataset was processed through an identical Salmon-based transcript quantification pipeline (utilizing read-length-matched k-mer sizes) and supplemented by five supporting analyses: (i) a housekeeping gene negative control to exclude pipeline artifacts and genomic DNA contamination, (ii) splice junction visualization via Sashimi plots to confirm *bona fide* splicing events, (iii) independent coding potential assessment (CPAT), (iv) NMD susceptibility prediction, and (v) *in silico* structural mapping of the predicted truncated protein to the canonical NKG2D domain architecture.

### 3.2 Retained-Intron *Klrk1* Isoforms Change with T Cell Activation and Differentiation Across Four Datasets

To assess whether *Klrk1* intron retention is a dynamic, activation-induced regulatory mechanism rather than a constitutive splicing artifact, it is first necessary to define its baseline expression in the quiescent state. To map this baseline, we quantified transcript-level expression in naive CD4⁺ (T.4.Nve; n=4) and naive CD8⁺ T cells (T.8.Nve; n=2) from unchallenged mice, together with naive P14 TCR-transgenic CD8⁺ cells (T8.TN.P14; n=2). Total *Klrk1* expression was minimal across both lineages (mean 0.76 TPM for naive CD4⁺; 15.0 TPM for naive CD8⁺; 10.6 TPM for naive P14 CD8⁺). Crucially, the retained-intron transcript *Klrk1-203* (ENSMUST00000137660), our primary isoform of interest, was below the detection limit (0.00 TPM) in every one of these eight resting samples. The canonical protein-coding isoform *Klrk1-201* (ENSMUST00000032252) was likewise undetectable in these naive cells.

Klrk1-203 was likewise undetectable in the naive P14 TCR-transgenic CD8⁺ samples. Across all eight resting samples examined here (four naive CD4⁺, two naive CD8⁺, and two naive P14 CD8⁺) Klrk1-203 was at or below the detection limit, providing a consistent baseline against which antigen-experienced populations can be compared.

We note one technical caveat in this dataset. In the naive samples, the canonical protein-coding isoform Klrk1-201 was also estimated at 0.00 TPM while total Klrk1 remained low but non-zero, a pattern that is difficult to reconcile biologically and most likely reflects the limited power of 25 bp reads to resolve isoform assignment at this locus when overall expression is low. We therefore treat the ImmGen naive values as evidence for the absence of an appreciable Klrk1-203 signal rather than as quantitative estimates, and anchor the quantitative claims of this study on the longer-read datasets (GSE119943, GSE147371, GSE203167).

Because NKG2D function is fundamentally linked to CD8⁺ T cell cytotoxicity, we next traced isoform kinetics across the antigen-experienced CD8⁺ populations in this compendium, all of which derive from an LCMV infection model. In terminal effector cells at LCMV day 7 (T8.TE; n=2), total Klrk1 expression rose to 311.7 TPM and Klrk1-203 to 49.6 TPM, accounting for 15.6% of the total Klrk1 pool. Memory precursor cells at the same time point (T8.MP; n=2) showed a comparable pattern (244.3 TPM total; Klrk1-203 31.3 TPM, 12.8%). At LCMV day 180, central memory cells (T8. Tcm; n=2) reached 478.9 TPM total with *Klrk1-203* at 90.6 TPM (18.9%), and a single effector memory sample (T8.Tem; n=1) reached 691.2 TPM total with *Klrk1-203* at 144.4 TPM, the highest proportion observed in this study at 20.9%.

Across these four antigen-experienced ImmGen populations, the Klrk1-203 fraction remained within a comparatively narrow band (12.8–20.9%), and the total retained-intron proportion likewise clustered between 21.7% and 34.4%. The single effector memory value should be interpreted with caution given n=1. Together, these data indicate that rather than functioning as a simple binary switch, Klrk1-203 usage operates as a graded feature of antigen-experienced CD8⁺ T cell states, present at appreciable levels across both effector and memory fates (Figure 4, Figure 5, Table 5). The age-matched, within-experiment gradient observed in the LCMV dataset described below (GSE119943, day 4.5 to day 7) addresses the differentiation axis under more controlled conditions.

**Figure 4.**
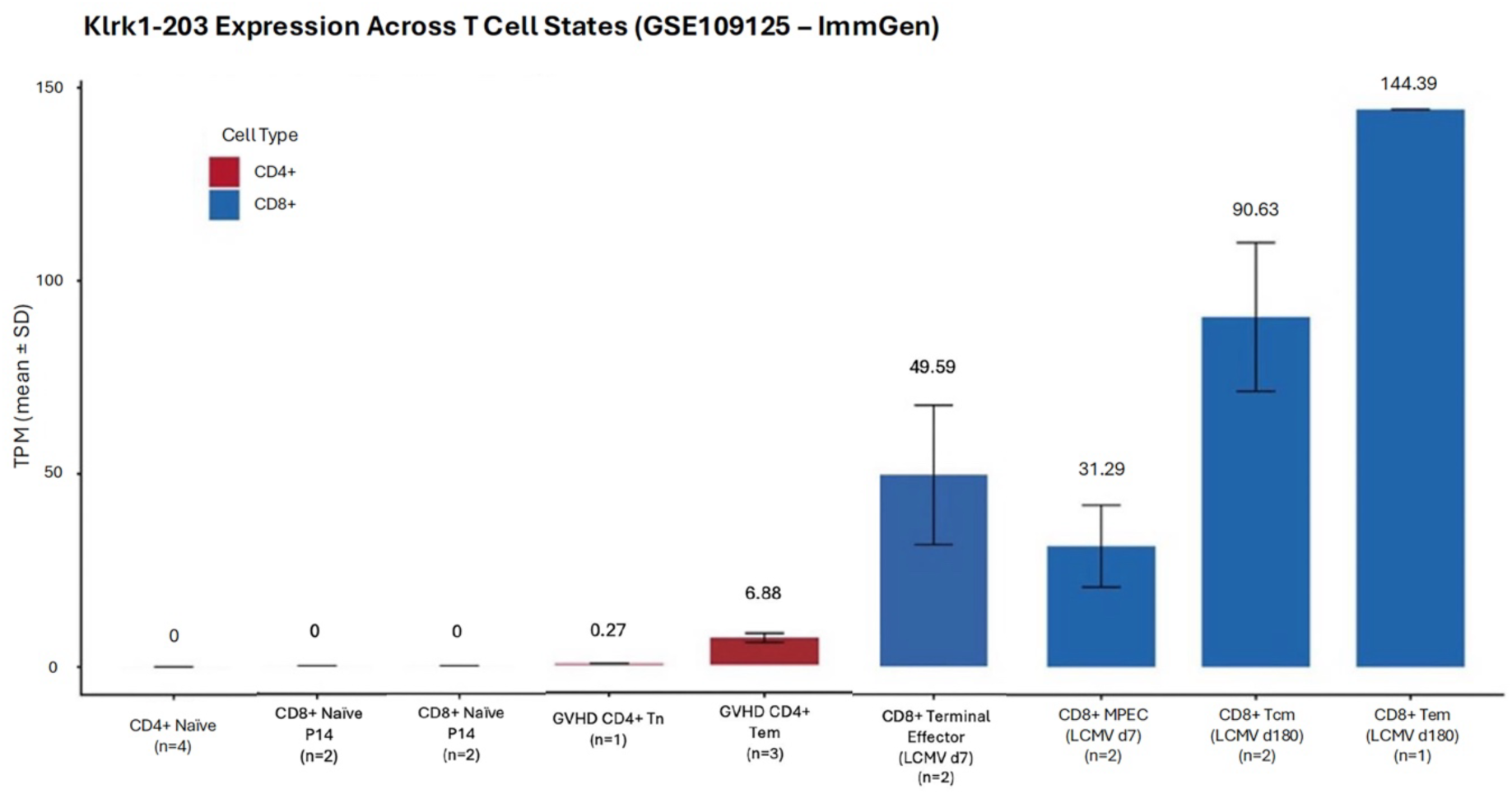
Klrk1-203 expression across T cell states. Absolute expression of the retained-intron transcript Klrk1-203 (ENSMUST00000137660) in transcripts per million (mean ± SD), across CD4⁺ and CD8⁺ T cell populations from two datasets. ImmGen (GSE109125) populations comprise naive CD4⁺ (n=4), naive CD8⁺ (n=2) and naive P14 TCR-transgenic CD8⁺ cells (n=2) from unchallenged mice, together with terminal effector (LCMV d7, n=2), memory precursor (LCMV d7, n=2), central memory (LCMV d180, n=2) and effector memory (LCMV d180, n=1) cells from an LCMV infection model. GVHD populations (GSE147371) comprise allogeneic CD4⁺ naive (Tn, n=1) and effector memory (Tem, n=3) cells at day 14 post-transplant. Klrk1-203 is undetectable in all eight unchallenged naive samples and is present at appreciable levels in every antigen-experienced population. *Klrk1-202* is reported within the combined *Klrk1-205/202* unit. Where n=1, no error bar is shown.

**Figure 5.**
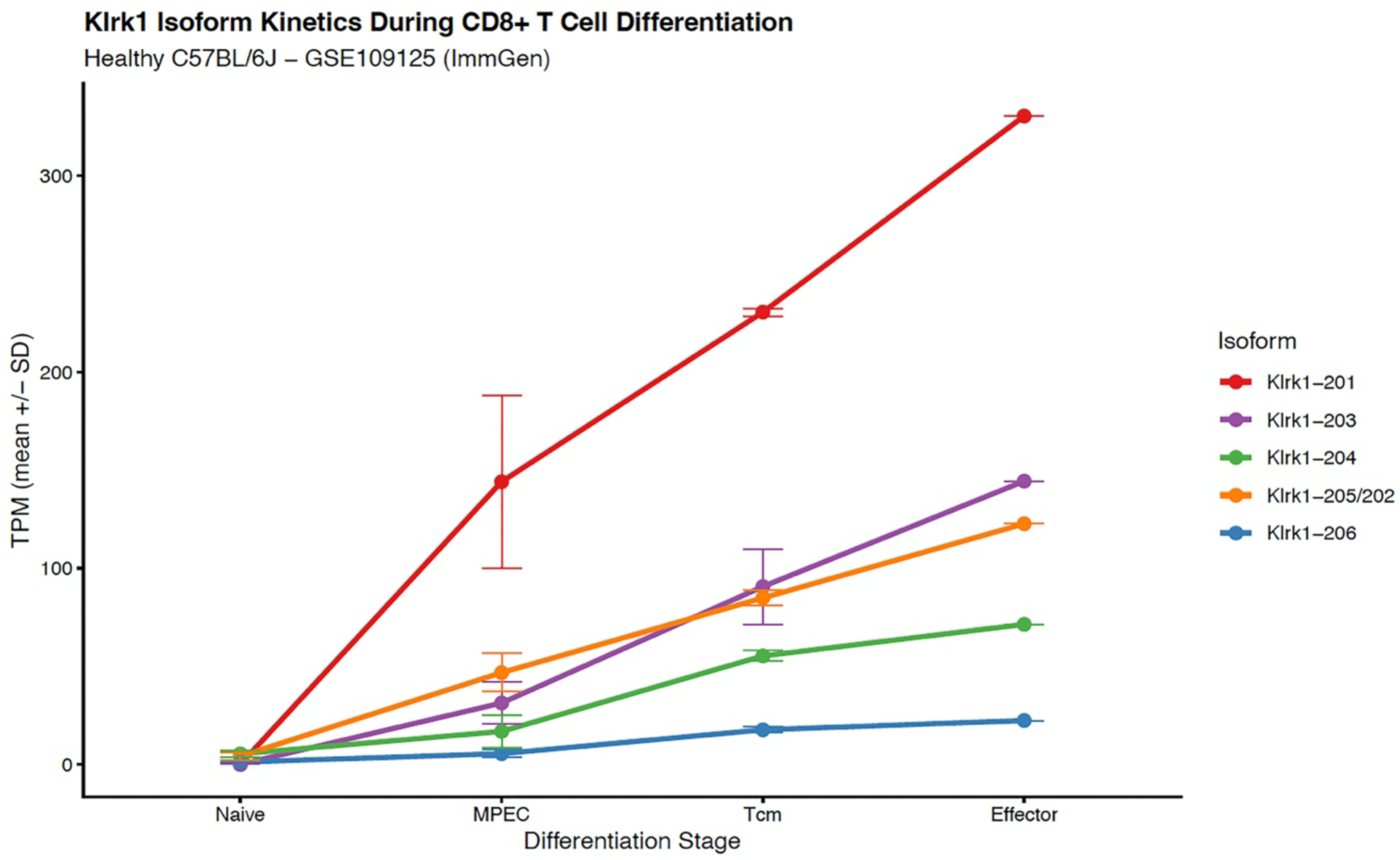
Klrk1 isoform kinetics across CD8⁺ T cell differentiation. Mean transcript-level expression (TPM ± SD) of each Klrk1 reporting unit across ImmGen CD8⁺ T cell populations (GSE109125): naive cells from unchallenged mice, and memory precursor (LCMV d7), central memory (LCMV d180) and effector memory (LCMV d180) cells from an LCMV infection model. All five reporting units rise with antigen experience, but Klrk1-203 increases from an undetectable baseline whereas Klrk1-204 and Klrk1-206 are already present in naive cells. Klrk1-202 is reported within the combined Klrk1-205/202 unit. Where n=1, no error bar is shown.

**Table 5.**
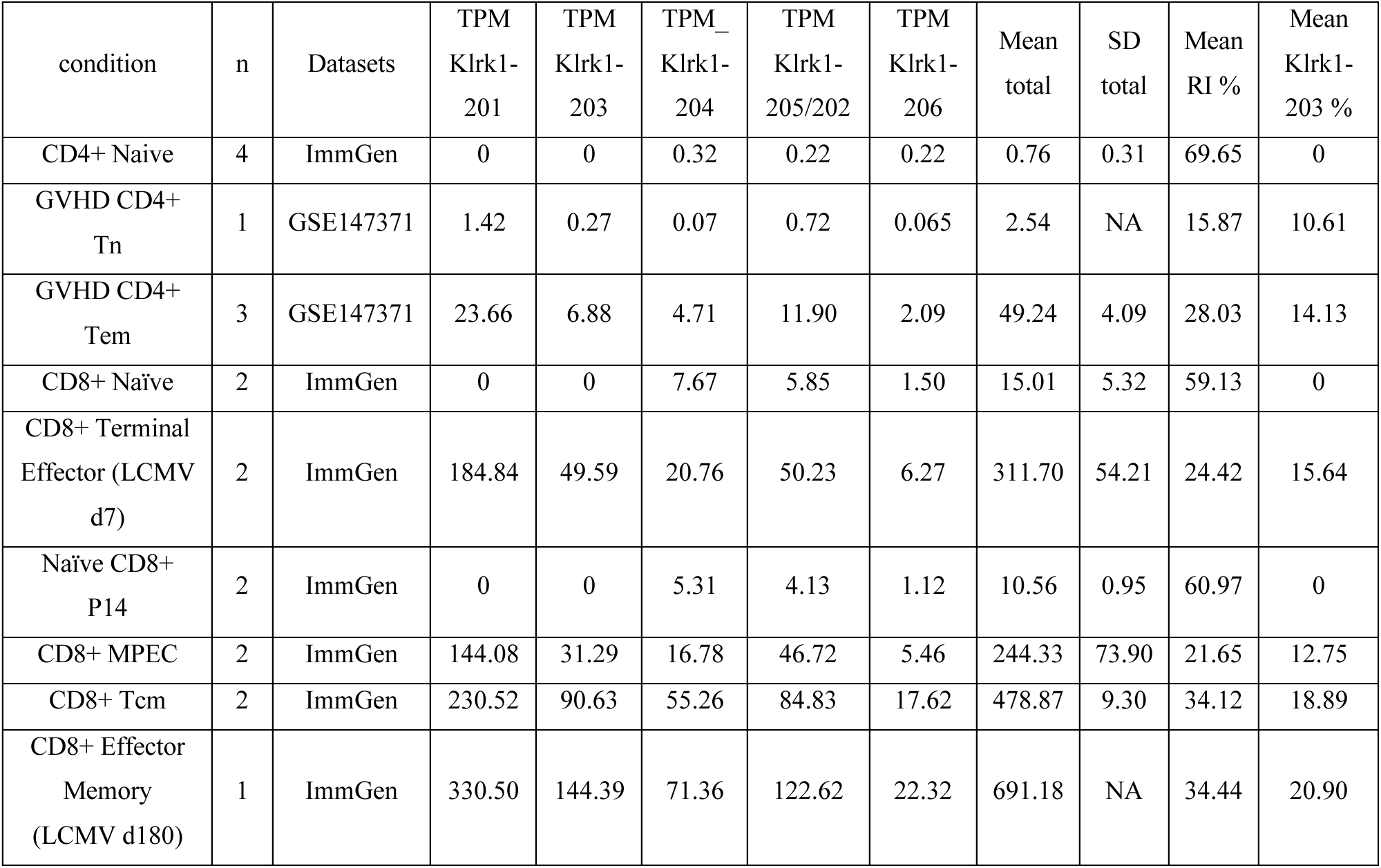
Master *Klrk1* isoform usage across T cell states in the ImmGen and GVHD datasets. Cross-dataset summary of mean transcript-per-million (TPM) expression levels for each *Klrk1* reporting unit. The total retained-intron (RI) percentage represents the sum of *Klrk1-203*, *-204*, and *-206* relative to total *Klrk1* TPM. Note the strict absence of *Klrk1-203* in naive CD4+ and CD8+ T cells (6-week), contrasting with its robust induction in antigen-experienced effector and memory subsets. Each row derives from a single dataset as indicated; values are not pooled across datasets. GSE119943 is reported separately in Table 6.

To determine whether this regulatory architecture is conserved across cell lineages and pathological contexts, we analyzed allogeneic CD4^+^ T cells from a murine GVHD model (GSE147371). Total *Klrk1* expression in GVHD CD4+ effector memory (Tem) cells (49.24 TPM) was approximately 14-fold lower than in CD8^+^ effector memory cells at LCMV day 180 (691.2 TPM). However, the proportional splicing architecture was remarkably conserved: retained-intron isoforms constituted 28% of total *Klrk1* transcription in GVHD CD4+ Tem cells, closely mirroring the 34.4% observed in ImmGen CD8+ effector memory cells. *Klrk1-203* alone contributed 14.1% of the total *Klrk1* pool in these GVHD Tem cells. The presence of a comparable proportion of retained-intron transcripts in a distinct cellular lineage and disease state suggests that this splicing pattern is neither confined to CD8+ cells nor strictly limited to healthy homeostatic conditions. While the precise percentages should be interpreted cautiously given the small sample sizes, the overarching conservation of this regulatory mechanism is clearly evident (Figure 6).

**Figure 6.**
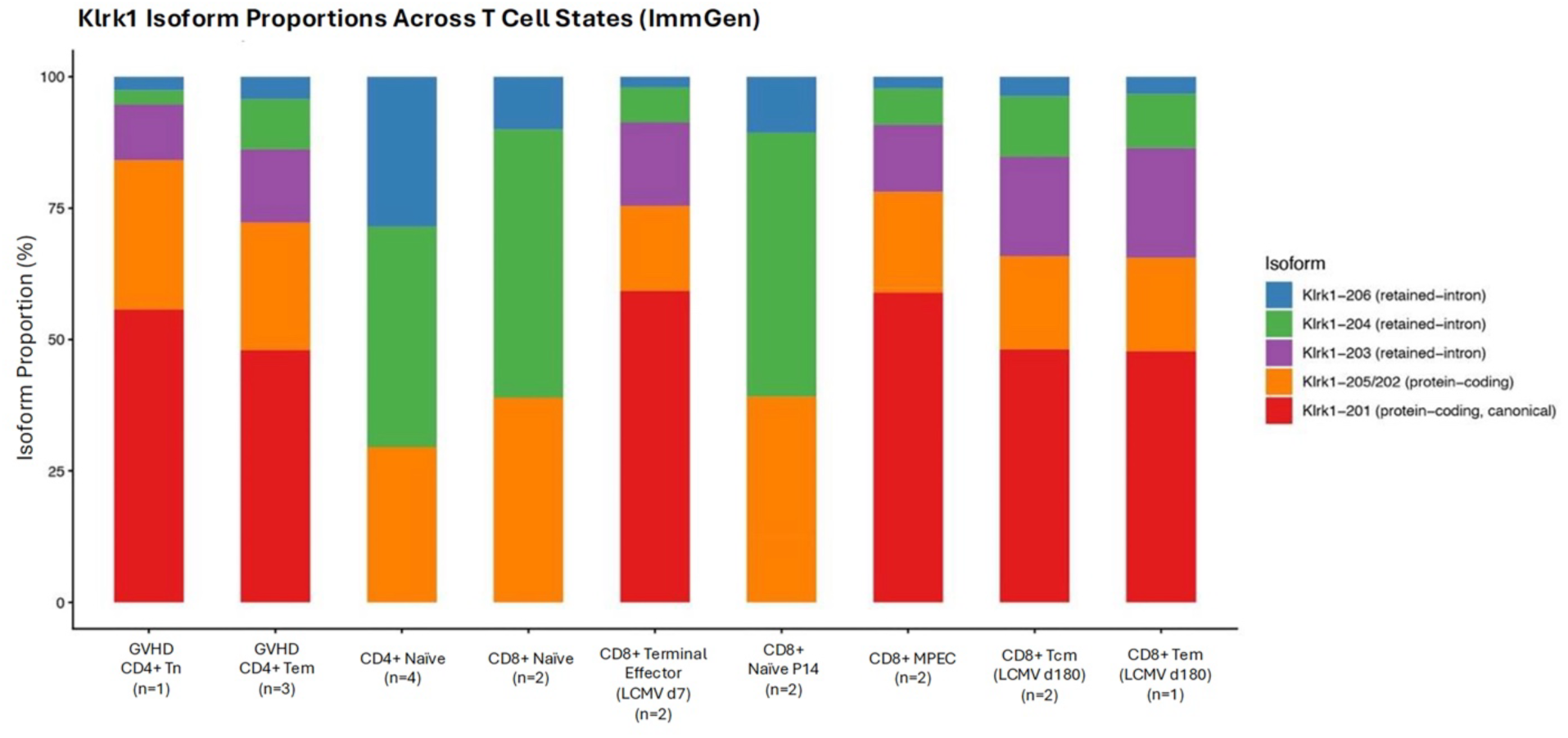
Klrk1 isoform proportions across T cell states. Stacked-bar visualisation of Klrk1 isoform composition, expressed as a percentage of the total Klrk1 transcript pool, across the same populations shown in Figure 4 (ImmGen GSE109125 and GVHD GSE147371). In unchallenged naive populations the retained-intron pool consists almost entirely of Klrk1-204 and Klrk1-206, with Klrk1-203 absent; in antigen-experienced populations Klrk1-203 emerges as a substantial fraction. The same pattern is seen in allogeneic GVHD CD4⁺ effector memory cells, where retained-intron isoforms constitute 28.0% of total Klrk1 and Klrk1-203 alone accounts for 14.1%, indicating that this splicing pattern is not restricted to the CD8⁺ lineage. Klrk1-202 is reported within the combined Klrk1-205/202 unit.

### 3.3 Klrk1-203 Induction Tracks with Effector and Memory Differentiation in an Independent LCMV Dataset (GSE119943)

While the prior datasets establish that *Klrk1* intron retention occurs across different lineages and pathological states, they are either cross-sectional or rely on severe inflammatory models. To test whether this splicing program is a fundamental, dynamic feature of a physiological immune response, we analyzed a fourth independent dataset (GSE119943) profiling P14 CD8^+^ T cells responding to acute (LCMV-Armstrong) and chronic (LCMV-Clone 13) viral infection. Unlike the cross-sectional ImmGen series, this dataset provides a robust within-experiment differentiation gradient.

In the acute infection model, *Klrk1-203* was below the detection limit (0.00 TPM) in early effector cells at day 4.5, a stage at which cells are highly activated and proliferating but not yet terminally differentiated (33, 35). At this time point, total *Klrk1* expression was heavily dominated by the protein-coding isoforms (*Klrk1-201*, 63%; *Klrk1-205/202*, 31% of transcripts), with a total retained-intron fraction of only 6.2%. By day 7, both short-lived effector cells (SLECs) and memory precursor effector cells (MPECs) had actively induced *Klrk1-203* to 19.5 ± 4.5 and 16.2 ± 4.0 TPM, respectively. This induction was accompanied by an approximately 2.9-fold increase in the total retained-intron fraction (reaching ∼18%) and a reciprocal decline in the canonical *Klrk1-201* fraction (from 63% to ∼57%). Thus, within a single, controlled experiment, *Klrk1-203* behaves specifically as a marker of effector/memory differentiation rather than merely a byproduct of initial T cell activation. Ultimately, these data indicate that Klrk1 intron retention is coupled to differentiation stage of T cell fate commitment, engaged comparably by both terminal-effector and memory-fated lineages (Figure 3; Table 6).

**Table 6.** *Klrk1* isoform expression dynamics during acute and chronic LCMV infection (GSE119943). Mean TPM values for *Klrk1* isoforms in P14 CD8+ T cells responding to acute (LCMV-Armstrong) or chronic (LCMV-Clone 13) viral infection. The retained-intron (RI) fraction represents the sum of all intron-retaining transcripts as a percentage of total *Klrk1*. *Klrk1-203* demonstrates distinct differentiation-coupled kinetics, remaining absent in early effectors (Day 4.5) before upregulating significantly in Day 7 SLEC and MPEC populations.

| Condition | n | GSM | Klrk1-201 | Klrk1-205/202 | Klrk1-204 | Klrk1-203 | Klrk1-206 | Total TPM | RI % | 203 % |
| --- | --- | --- | --- | --- | --- | --- | --- | --- | --- | --- |
| Arm D4.5 Early Effector | 3 | GSM3568595, 96, 97 | 89.8 | 44.3 | 7.2 | 0.0 | 1.7 | 142.9 | 6.2 | 0.0 |
| Arm D7 MPEC | 3 | GSM3568614, 15, 16 | 140.8 | 63.0 | 25.7 | 16.2 | 2.6 | 248.2 | 17.9 | 6.5 |
| Arm D7 SLEC | 3 | GSM3568611, 12, 13 | 160.3 | 68.3 | 26.8 | 19.5 | 2.8 | 277.6 | 17.7 | 7.0 |
| CI13 D7 Progenitor-like | 2 | GSM3568609, 10 | 55.2 | 34.3 | 8.9 | 2.7 | 2.4 | 103.5 | 12.9 | 2.6 |
| CI13 D7 Terminally Exhausted | 2 | GSM3568607, 08 | 62.1 | 45.4 | 10.5 | 3.2 | 3.1 | 124.3 | 13.3 | 2.6 |
| pMIG CI13 Progenitor-like | 3 | GSM3388395, 96, 97 | 48.2 | 71.9 | 10.1 | 10.9 | 3.2 | 144.3 | 16.7 | 7.5 |
| pMIG CI13 Terminally Exhausted | 3 | GSM3388398, 99, 3388400 | 71.5 | 99.6 | 20.2 | 6.3 | 4.3 | 201.9 | 15.1 | 3.0 |

To determine how prolonged antigen stimulation and the onset of T cell exhaustion impact this splicing program, we next evaluated the chronic infection cohort. In chronic infection, where CD8^+^ T cells follow an exhaustion rather than a canonical effector/memory trajectory, *Klrk1-203* was induced more modestly (*TCF1*+ progenitor-exhausted, 2.7 TPM; terminally exhausted, 3.2 TPM; overall retained-intron fraction, ∼13%). The empty-vector (pMIG) control cells showed higher Klrk1-203 usage than their non-transduced counterparts (progenitor-like, 10.9 TPM; ∼7.5% of total Klrk1). These cells were activated in vitro, retrovirally transduced and sorted before adoptive transfer, and are therefore not directly comparable to the non-transduced infection groups; they also showed a markedly higher Klrk1-205/202 fraction (∼50% versus 33–36%), consistent with in vitro activation altering the isoform landscape at this locus. We report them for completeness but exclude them from the differentiation-axis interpretation. However, these chronic and transduced groups (n=2–3) exhibited high sample-to-sample variability, with per-sample *Klrk1-203* fractions spanning roughly 0% to 14% within a single group. Consequently, we treat this chronic exhaustion data as exploratory rather than a basis for definitive conclusions.

Interestingly, across all seven GSE119943 conditions, the canonical *Klrk1-201* isoform constituted a major fraction of the transcript pool (33–63%) but was not always the most abundant. In the pMIG-transduced cells, the alternative-promoter protein-coding unit *Klrk1-205/202* (∼50%) exceeded it, and *Klrk1-205/202* remained a substantial fraction throughout the entire dataset (25% in acute effectors, ∼31% at day 4.5, and 33–50% in chronic and transduced cells). Together, these data indicate that intron retention is not an isolated, all-or-none switch, but rather one specific arm of a much broader, differentiation-coupled remodeling of the entire *Klrk1* isoform landscape.

To assess whether this splicing behavior is a targeted regulatory event rather than a generalized artifact of transcriptional noise or incomplete processing, we evaluated housekeeping negative controls within this dataset. Within this dataset, the retained-intron fraction at *Klrk1* (median 16.2%) robustly exceeded that of *Actb* (0.83%) and *Gapdh* (2.28%) by approximately 20-fold and 7-fold, respectively. Furthermore, Sashimi visualization of representative acute-effector (SLEC) (35) and chronic-exhausted (36) samples confirmed *bona fide* splice-junction usage across the *Klrk1* locus, with junction-spanning read counts scaling proportionally with total *Klrk1* expression (1,299 versus 744 junction-spanning reads). Although the absolute magnitude of the retained-intron fraction was lower in GSE119943 (∼18% at the effector peak) compared to the ImmGen effector memory population (34%), an expected variance given the cross-dataset differences in sequencing platforms, read lengths, and sorting strategies, the fundamental qualitative pattern was reproduced in full. Ultimately, *Klrk1-203* is undetectable in the least-differentiated cells and is induced specifically upon effector and memory differentiation, consistent with intron retention being a regulated feature of T cell maturation

Because the variant transcripts *Klrk1-203*, *Klrk1-204*, and *Klrk1-206* retain distinct intronic segments and may be governed by independent regulatory mechanisms, we sought to deconvolve the total retained-intron fraction into its constituent parts across all conditions (Table S1). (As detailed in the Methods, *Klrk1-202* is reported within the combined *Klrk1-205/202* unit). Crucially, this high-resolution analysis revealed that the differentiation-associated signal tracked specifically with *Klrk1-203* rather than with the retained-intron fraction as a whole. While *Klrk1-203* comprised 0% of the total *Klrk1* pool in healthy naive populations and expanded to 12.8–20.9% across the ImmGen effector and memory subsets, the total retained-intron fraction was paradoxically highest in naive cells (e.g., ∼70% in naive CD4+ T cells). In these resting cells, the retained-intron pool consisted almost entirely of *Klrk1-204* and *Klrk1-206*, with *Klrk1-203* contributing nothing. One naive-phenotype sample deviated from this baseline: the allogeneic GVHD donor naive (Tn) sample exhibited Klrk1-203 at ∼11%. This sample is a single replicate drawn from an allogeneic transplant recipient rather than an unchallenged animal, and its total Klrk1 expression was only 2.5 TPM, so its isoform proportions rest on very few reads. We report it as observed rather than excluding it. Across the eight unchallenged naive samples, Klrk1-203 was uniformly below the limit of detection. Ultimately, these data demonstrate that the total retained-intron fraction is too blunt an instrument to serve as a clean readout of differentiation state. The highly informative, activation-specific metric is the isolated induction of *Klrk1-203* with the necessary bioinformatic caveat that distinguishing it from the co-retained *Klrk1-204* transcript inherently relies on model-based quantification.

### 3.4 Total *Klrk1* Rises While the Retained-Intron Pool Is Remodeled After Transplantation (GSE203167)

Having established *Klrk1-203* induction in steady-state differentiation and viral infection, we next sought to determine whether this splicing program is engaged during the extreme, clinically relevant immunological stress of allogeneic transplantation, and how it behaves in the absence of a canonical memory regulator. To this end, we analyzed the GSE203167 dataset (n=12; 4 conditions × 3 biological replicates), which compares wild-type (WT) and *TCF-7* conditional knockout (cKO) CD8+ T cells before and after allogeneic transplantation. Total *Klrk1* expression increased substantially post-transplant: in WT cells, the mean total *Klrk1* TPM rose from 35.2 (Pre-Tx) to 72.6 (Post-Tx D7). In *TCF-7* cKO cells, this increase was even more pronounced, surging from 37.9 (Pre-Tx) to 276.7 (Post-Tx D7), an approximate 7-fold induction (Figure 7A).

**Figure 7.**
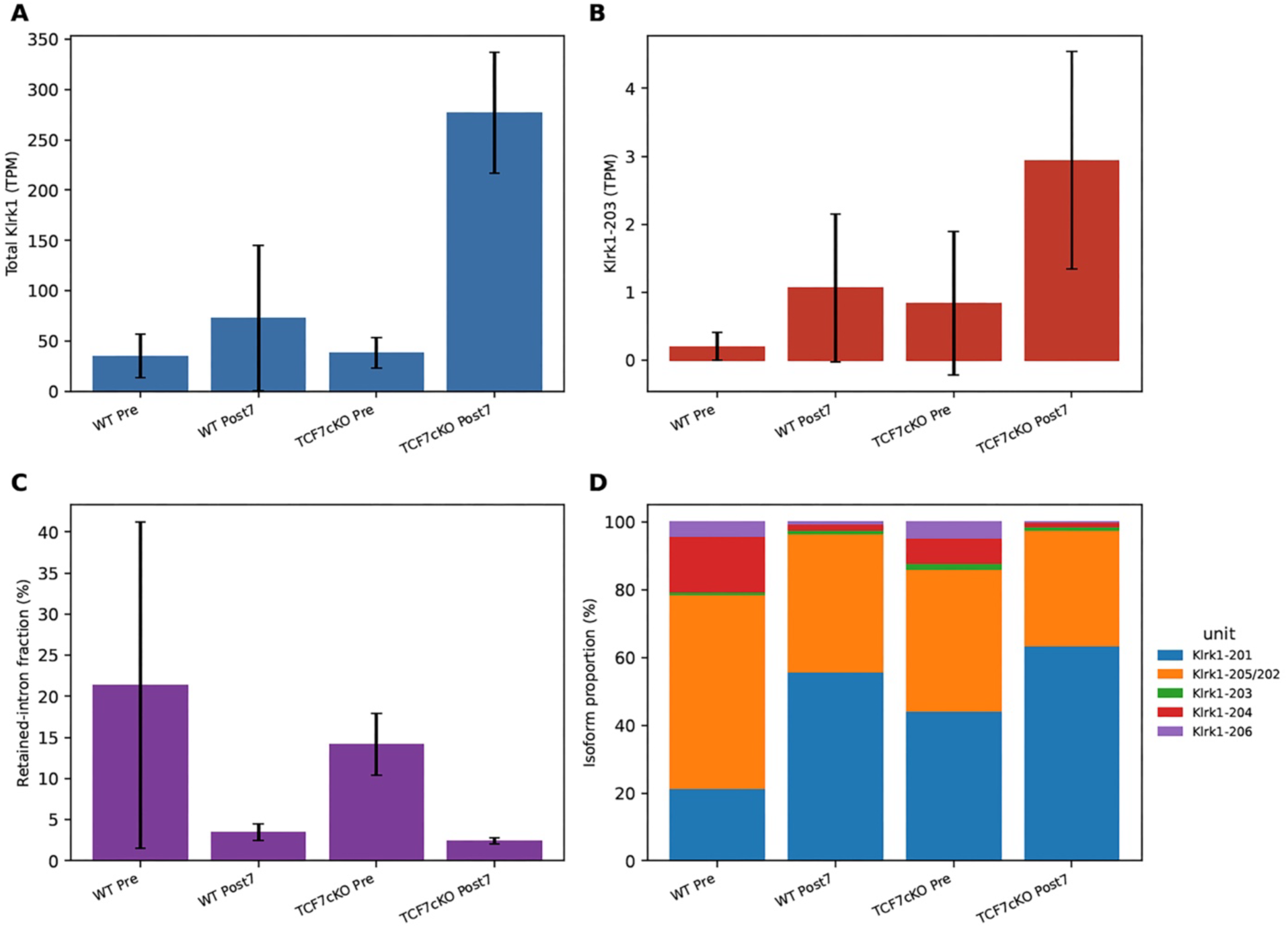
*Klrk1* isoform usage and remodeling following allogeneic transplantation. Transcriptomic profiling of wild-type (WT) and TCF-7 conditional knockout (cKO) CD8^+^ T cells pre-transplant and on day 7 post-transplant (GSE203167, n=3 per condition). **(A)** Absolute expression of total *Klrk1* (TPM, mean ± SD) showing massive induction post-transplant. **(B)** Absolute expression of *Klrk1-203* (TPM, mean ± SD), demonstrating activation-induced upregulation. **(C)** The total retained-intron fraction (*Klrk1-203* + *204* + *206* as % of total *Klrk1*) drops sharply post-transplant (WT 21% → 3%; TCF-7 cKO 14% → 2%). **(D)** Detailed isoform proportions reveal this fractional drop is driven by the post-transplant collapse of the baseline *Klrk1-204* transcript and the profound induction of canonical isoforms, particularly in TCF-7 cKO cells.

Crucially, absolute expression of the retained-intron transcript *Klrk1-203* (ENSMUST00000137660) also increased post-transplant across both genotypes. *Klrk1-203* levels rose from 0.21 to 1.07 TPM in WT cells (∼5-fold induction) and from 0.84 to 2.93 TPM in *TCF-7* cKO cells (∼3.5-fold induction) (Figure 7B). These data confirm that absolute *Klrk1-203* transcription is robustly induced upon post-transplant activation, even from a relatively low pre-transplant baseline. However, as detailed below, the proportional remodeling of the transcript pool in this specific allo-transplant setting presents a more complex picture than the straightforward induction observed in the ImmGen and GVHD models.

When we examined the relative isoform proportions in this dataset, a nuanced and seemingly paradoxical observation emerged. Although absolute *Klrk1-203* expression robustly rose post-transplant, the overall retained-intron fraction decreased in both genotypes (WT: 21% Pre-Tx → 3% Post-Tx; *TCF-7* cKO: 14% Pre-Tx → 2% Post-Tx) (Figure 7C). This apparent inversion (relative to the naive-to-memory trajectories observed in the ImmGen and GVHD datasets) is explained by the independent regulatory behavior of the other retained-intron isoforms, principally *Klrk1-204*.

Before transplantation, the retained-intron pool was heavily dominated by *Klrk1-204* (accounting for roughly 16% of total *Klrk1* in WT Pre-Tx cells). Following transplantation, *Klrk1-204* expression precipitously collapsed (falling to roughly 2%), whereas *Klrk1-203* rose modestly as a proportional fraction of the overall Klrk1 pool (from ∼0.6% to ∼1.5%) while simultaneously rising in absolute expression. As a result, *Klrk1-203* comprised a rapidly growing share of a shrinking retained-intron pool (shifting from ∼4% to ∼29% of all retained-intron transcripts – average per sample-in WT cells). Thus, the precipitous fall in the total retained-intron fraction reflects the selective collapse of *Klrk1-204* rather than a failure to induce *Klrk1-203* (Figure 7D). Ultimately, this finding reveals that the intron-retention program is not a monolithic entity; individual retained-intron isoforms operate concurrently with, but at distinct kinetics from, canonical isoform induction, and this balance is highly context-dependent. Furthermore, the *TCF-7* cKO genotype exhibited the most pronounced canonical-isoform dominance post-transplant, suggesting that this key memory transcription factor may actively govern the balance of alternative splicing during severe immunological stress.

### 3.5 The Retained-Intron Signal Is Gene-Specific and Detectable at the Read Level

A critical concern in retained-intron analyses is whether the observed signal reflects *bona fide* intron retention or is instead attributable to technical artifacts such as genomic DNA contamination, incomplete library size selection, or quantification bias. To address this, we performed an identical isoform proportion analysis on two highly expressed, constitutively spliced housekeeping genes, *Actb* and *Gapdh*, across all 50 samples of the four independent datasets (GSE147371, GSE109125, GSE203167, and GSE119943).

The retained-intron fraction at the *Klrk1* locus (median 17%, interquartile range 12–31%) was approximately 42-fold higher than at *Actb* (median 0.40%, range 0–1%) and 9-fold higher than at *Gapdh* (median 1.87%, with a single outlier at 28% occurring in a sample with exceptionally low total *Gapdh* TPM). Both housekeeping genes exhibited retained-intron proportions entirely consistent with baseline background transcriptional noise (Figure 1). If the *Klrk1* retained-intron signal were attributable to systemic genomic DNA contamination or pipeline-level artifacts, similar proportional elevations would be expected at *Actb* and *Gapdh*; however, they were completely absent. This contrast indicates that the *Klrk1* retained-intron program is a targeted, gene-specific biological phenomenon rather than a technical artifact.

A further alternative explanation is that the apparent Klrk1-203 signal simply tracks total Klrk1 abundance, appearing only once the locus is sufficiently expressed for the quantification model to apportion reads to a minor isoform. The GSE119943 data argue against this. Day-4.5 early effector cells expressed 142.9 TPM total Klrk1 with no detectable Klrk1-203, whereas Cl13 progenitor-exhausted cells expressed 103.5 TPM with 1.8%, and day-7 effector populations reached 6.5–7.0%. Total Klrk1 abundance alone therefore does not predict the retained-intron signal, which instead tracks differentiation state.

Furthermore, if the elevated intron-4 signal reflected generalized nuclear RNA or genomic DNA contamination rather than actively regulated alternative splicing, all retained-intron isoforms of *Klrk1* would be expected to rise in tandem. Crucially, they do not. *Klrk1-203* is induced highly specifically, remaining undetectable (0%) in every naive population before rising to 13–21% in ImmGen effector and memory cells, while *Klrk1-204* and *Klrk1-206* follow distinct and often opposing transcriptional trajectories. This strict isoform selectivity argues against a blanket contamination artifact and is consistent with *Klrk1-203* induction is driven by active post-transcriptional regulation.

### 3.6 Sashimi Plot Confirms Splice Junction Usage at the Klrk1 Locus

To provide direct, alignment-level evidence of splice junction usage and to visualize the retained-intron events at the *Klrk1* locus, we generated Sashimi plots from HISAT2-aligned BAM files of two representative GSE203167 samples (one WT Pre-Tx and one WT Post-Tx D7). Visual inspection of the coverage profiles confirmed canonical splice junctions consistent with the protein-coding *Klrk1-201* and *Klrk1-205/202* transcripts in both conditions. Coverage across the Klrk1 locus was visibly higher in the Post-Tx D7 sample than in the Pre-Tx baseline, consistent with the activation-induced increase in Klrk1 expression seen at the transcript level (Figure 2). Crucially, coverage was also distinctly observed across the intronic regions corresponding to the *Klrk1-203* retained intron, directly matching our Salmon-based quantification.

While Sashimi plot visualization confirmed *bona fide* splicing activity across the *Klrk1* locus, we sought to quantitatively verify junction usage. To address this, we extracted and counted the splice junction reads directly from these HISAT2 alignments. All four annotated canonical *Klrk1* junctions were robustly detected in both conditions, indicating that the baseline splicing machinery is fully active at this locus and that intron retention is not a mere consequence of global splicing failure. Specifically, the exon 4–exon 5 junction (the canonical splicing event whose execution removes intron 4) was supported by 147 reads in the WT Pre-Tx sample and 202 reads in the WT Post-Tx D7 sample. This 37% increase in canonical junction usage confirms that *Klrk1-203* induction occurs concurrently with active, successful splicing of the primary transcript, rather than from a generalized collapse of spliceosome function.

Junction-based percent intron retention (PIR) at intron 4 (computed as retention-supporting reads divided by the sum of retention and canonical exon 4–exon 5 junction reads) was 2.0% in WT Pre-Tx cells and 5.2% in WT Post-Tx D7 cells. Because sequencing reads crossing the exon 4–intron 4 boundary cannot be uniquely assigned to a single transcript, this read-level PIR value measures total intron-4 retention rather than *Klrk1-203* specifically. Consequently, it runs slightly above the Salmon-derived *Klrk1-203* transcript fraction in these same samples (0.7–1.1%), a discrepancy entirely consistent with the co-expressed *Klrk1-204* isoform also contributing boundary-spanning reads. Both measures confirm that intron-4 retention constitutes only a minor fraction of the total *Klrk1* pool in this acute post-transplant dataset, which is overwhelmingly dominated by canonical isoforms. By contrast, the retained-intron program is much more active in the ImmGen effector/memory and GSE119943 day-7 effector populations, where *Klrk1-203* reaches 7–21% of total *Klrk1* expression.

To rigorously confirm the transcript-level *Klrk1-203* signal at the raw read level across the remaining datasets, we evaluated PIR utilizing the same junction-spanning logic. In the GVHD CD4+ dataset (GSE147371, 75 bp reads), all four samples showed the expected pattern: the three-effector memory (Tem) samples carried abundant spliced reads (49–110) with an intron-4 PIR of 14–23%, highly concordant with their Salmon-derived *Klrk1-203* fractions (10–16%). Meanwhile, the naive (Tn) sample yielded almost no signal (3 spliced reads), precisely as expected for a resting cell state where *Klrk1* is minimally expressed (Table 7).

**Table 7.**
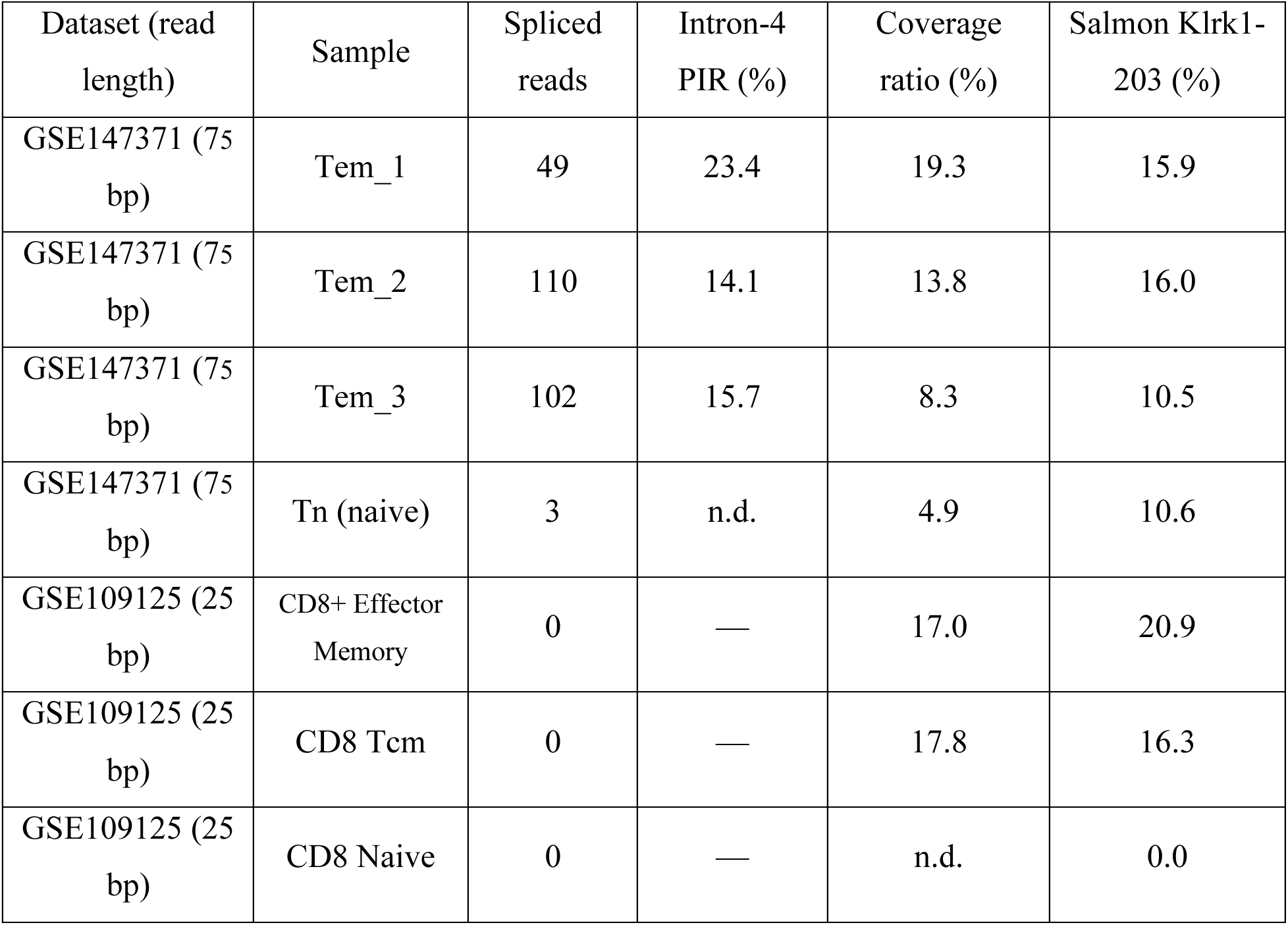
Alignment-level Percent Intron Retention (PIR) confirmation of intron-4 retention. Read-level validation of intron-4 retention derived directly from HISAT2-aligned BAM files, independent of Salmon transcript modeling. PIR is calculated as the ratio of reads spanning the exon 4–intron 4 boundary to the sum of retention and canonical (exon 4–exon 5) spliced reads. Read-length limitations in the 25 bp ImmGen dataset preclude standard junction-spanning PIR calculations; a read-length-independent coverage ratio is provided instead. Values are from single representative samples and therefore differ slightly from the group means reported in Table 5.

For the ImmGen dataset (GSE109125, 25 bp), the sequencing reads were too short to be reliably scored as junction-spanning which was a technical read-length limitation rather than a biological absence. We therefore utilized a read-length-independent intron-4 coverage ratio. This alternative metric closely tracked the Salmon estimates in the highly expressed effector/memory populations (17–18% coverage versus 16–21% Salmon fraction) and predictably dropped to near zero in unexpressed naive cells. Because the exon 4/intron 4 boundary is identically crossed by both *Klrk1-203* and the co-expressed *Klrk1-204*, these read-level metrics report overall intron-4 retention; the specific attribution to *Klrk1-203* relies on the Salmon quantification and our 203-versus-204/206 decomposition. Ultimately, the tight convergence between an isoform-inference-free read-level measure and our transcript-level model strongly argues that the quantified intron-4 retention is a genuine biological event rather than an artifact of short-read isoform ambiguity (Table 7).

### 3.7 Klrk1-203 Is Predicted to Escape NMD and Encode a Truncated, NKG2D-Related Product

To independently verify the non-coding annotation of *Klrk1-203* without relying solely on the Ensembl database, we applied the Coding Potential Assessment Tool (CPAT) to the ENSMUST00000137660 cDNA sequence using its mouse-specific logit model. CPAT yielded a coding probability score of 0.158 (well below the established mouse-specific coding cutoff of 0.44) and algorithmically classified Klrk1-203 as non-coding under the mouse-specific model, providing sequence-based support for, but not experimental confirmation of, the Ensembl retained-intron/non-coding annotation.

However, a central question for any retained-intron transcript is whether it is rapidly targeted for nonsense-mediated decay (NMD) or if it escapes this surveillance pathway to accumulate stably in the cell. Because the biological implications of *Klrk1-203* differ drastically under each scenario, we performed two complementary sequence analyses. First, we mapped the canonical *Klrk1-201* start codon onto the *Klrk1-203* transcript sequence; second, we scanned the full transcript in all three reading frames to identify the longest unbiased open reading frame (ORF).

Both approaches converged on the identical start codon. The canonical *Klrk1-201* ATG is retained at transcript position 112 within *Klrk1-203*, and the longest ORF identified by three-frame scanning initiates at this exact position in Frame +1. Translation from this ATG proceeds normally through the upstream exons before reading into the retained intron 4, ultimately terminating at transcript positions 475 to 477 (TGA) within the retained intron itself. This yields a predicted 121-amino-acid product (∼13.3 kDa). Critically, the TGA termination codon lies within the terminal exon, beginning at transcript position 475, approximately 180 nucleotides into exon 4 and leaving only 24 nucleotides of sequence downstream. The final exon-exon junction, EEJ3, occurs between transcript positions 295 and 296, and no exon-exon junction remains downstream of the termination codon. Thus, rather than being evaluated by its distance upstream of a downstream junction, this ORF satisfies the last-exon rule and is predicted to evade classical EJC-dependent NMD. This positional prediction does not establish complete resistance to all RNA surveillance pathways, as retained-intron transcripts may additionally be subject to nuclear retention, nuclear RNA degradation, inefficient export, or EJC-independent decay mechanisms.

The Kozak sequence context at position 112 also remains a favorable Kozak context (A at −3, G at +4), consistent with efficient translation initiation. These independent analyses identify the same ATG as the dominant candidate initiation site, providing a parsimonious primary translation model. Ultimately, these findings raise the intriguing possibility that *Klrk1-203* not only accumulates as a stable regulatory transcript, but may also be translated into a novel, truncated NKG2D-derived protein, which needs further experimental confirmation.

### 3.8 *Klrk1-203* Is Predicted to Encode a Truncated Protein Analogous to Human NKG2D-TR

To assess the potential functional consequences of Klrk1-203 translation, we mapped the predicted 121-amino-acid product onto the canonical murine NKG2D domain topology represented by UniProtKB O54709-1. UniProt annotates the canonical 232-amino-acid mouse NKG2D protein as comprising a cytoplasmic region at amino acids 1–66, a transmembrane helix at amino acids 67– 89, and an extracellular region at amino acids 90–232, with the C-type lectin domain spanning amino acids 122–228. Translation of Klrk1-203 initiates from the canonical Klrk1-201 ATG and preserves the first 96 amino acids of canonical NKG2D. The predicted product therefore retains the complete cytoplasmic region and transmembrane helix, followed by the first seven amino acids of the canonical extracellular region. Translation then diverges from the canonical Klrk1-201 sequence within the terminal retained-intron-associated region, producing an approximately 25-residue isoform-specific C-terminal sequence before termination. Consequently, the predicted Klrk1-203 product lacks the entire annotated C-type lectin domain of canonical mouse NKG2D.

This predicted domain architecture shows a notable structural parallel to human NKG2D-TR, a dominant-negative splice variant generated through retention of intron 4 in the human KLRK1 locus. Human NKG2D-TR retains the cytoplasmic and transmembrane regions but lacks the extracellular ligand-binding domain; experimental studies demonstrated that it associates with both DAP10 and full-length NKG2D, interferes with full-length NKG2D–DAP10 complex formation, and reduces functional NKG2D surface expression. Retention of the canonical transmembrane segment, including the conserved transmembrane arginine required for NKG2D– DAP10 assembly, suggests that the predicted Klrk1-203 product may similarly retain the capacity to associate with DAP10. However, this interaction, as well as the membrane insertion, stability, localization, and potential dominant-negative activity of the predicted murine product, requires direct biochemical validation. Together with its differentiation-coupled induction, reaching up to 144 TPM in CD8+ effector memory cells, favorable Kozak context, and predicted evasion of classical EJC-dependent NMD, these features support a testable model in which murine T cells may employ a structurally analogous post-transcriptional mechanism to restrain NKG2D signaling during peak activation. We emphasize, however, that the existence and function of the *Klrk1-203*-derived protein remain computational predictions until stable protein expression, subcellular localization, interaction with DAP10 or full-length NKG2D, and effects on NKG2D signaling are experimentally demonstrated.

### 3.9 A Proximal NFAT Motif in the Shared *Klrk1* Promoter

To determine whether the activation-induced transcription of *Klrk1* might be directly driven by canonical T cell receptor (TCR) signaling pathways, we scanned the shared *Klrk1* proximal promoter for Nuclear Factor of Activated T cells (NFAT) binding motifs. A perfect-consensus NFAT element (AATGGAAA; relative score 1.00) was identified 134 bp upstream of the canonical *Klrk1* transcription start site (TSS; motif 5′ edge at chr6:129,599,866–129,599,873). This element was confidently recognized by both *Nfatc1* and *Nfatc2* position weight matrices at high stringency (p < 1 × 10⁻⁴) (Figure 8).

**Figure 8.**
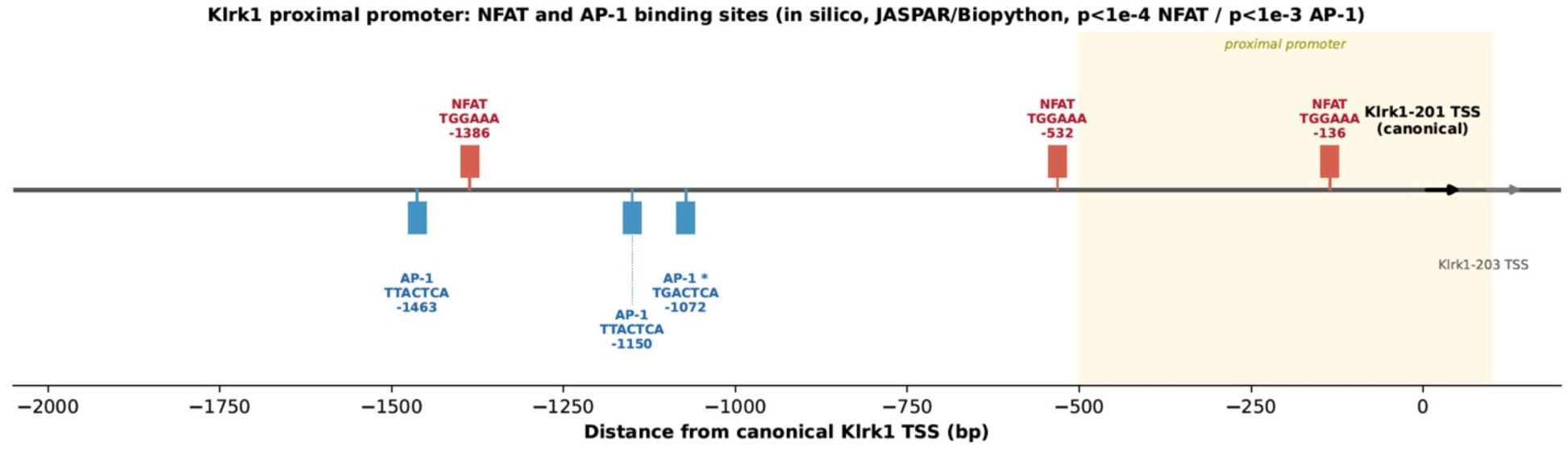
In silico NFAT and AP-1 motif map of the shared Klrk1 proximal promoter. Position weight matrix scan of a 2.3-kb window spanning −2000 to +288 bp relative to the canonical Klrk1-201 transcription start site (chr6:129,599,447–129,601,735, GRCm39), performed on both strands using JASPAR 2024 CORE (vertebrates) matrices and Biopython log-odds PSSMs. NFAT matches (red, above the axis) were called at a false-positive rate of 1 × 10⁻⁴; AP-1 matches (blue, below the axis) at the more permissive threshold of 1 × 10⁻³ and should be interpreted with corresponding caution. A perfect-consensus NFAT element lies within the proximal promoter, with two further NFAT core motifs at approximately −532 and −1386 bp. A consensus AP-1/TRE element (TGACTCA) is present at approximately −1072 bp alongside two lower-scoring AP-1 motifs. No NFAT:AP-1 composite elements were detected at an edge-to-edge spacing of ≤20 bp. The shaded region marks the proximal promoter; the Klrk1-203 transcription start site lies 88 bp downstream of the canonical Klrk1-201 TSS, so the two transcripts share this promoter. All matches are computational predictions and are presented as candidate regulatory elements rather than functionally validated sites.

Two additional NFAT core motifs (GGAAA) were located further upstream at positions −532 bp and −1386 bp. While a perfect AP-1/TRE element (TGACTCA) was present at −1072 bp alongside two lower-scoring AP-1 motifs, no tightly spaced NFAT:AP-1 composite elements were detected within 20 bp. Because a 2.3-kb promoter scanned on both strands is statistically expected to yield a few short motifs match by chance, and because this scan lacked a shuffled-sequence background model or cross-species conservation analysis, we present this as a candidate regulatory element rather than a functionally validated one. Furthermore, the AP-1 matches were called at a more permissive threshold (p < 1 × 10⁻³) and should be interpreted with caution.

Nevertheless, because *Klrk1-203* shares this primary proximal promoter with the canonical protein-coding transcript, an inducible, TCR-coupled NFAT input would theoretically drive the initial transcription of both pre-mRNAs simultaneously. This shared initiation architecture suggests that differential splicing and isoform-specific RNA stability may contribute substantially to the distinct expression kinetics. Experimental validation will ultimately be required to confirm this regulatory model.

## 4 Discussion

The activating immune receptor NKG2D is a critical driver of cytotoxic responses in both tumor immunity and alloimmune pathologies such as GVHD (8, 37). In humans, this potent signaling is tightly restrained post-transcriptionally by NKG2D-TR, a dominant-negative splice variant (18). However, the existence of a functionally analogous “brake” in the mouse (the foundational preclinical model for NKG2D-targeted therapies) has remained a critical missing piece of the regulatory puzzle. In this study, we offer a systematic, multi-dataset resolution of the murine *Klrk1* (NKG2D) transcriptomic landscape. Across 50 independent biological samples, we demonstrate that the murine *Klrk1* locus harbors a highly dynamic, activation-coupled retained-intron program. Specifically, the *Klrk1-203* isoform is not a fixed or constitutive transcript; it is below detection limit in naive, and early-effector T cells, yet robustly induced upon effector and memory differentiation (22, 38, 39). Crucially, *in silico* structural mapping localizes the retained intron of *Klrk1-203* exactly at the junction between the transmembrane and ligand-binding domains, mirroring the precise topological architecture of human NKG2D-TR (18).

Taken together, these computational observations nominate Klrk1-203 as a candidate for a post-transcriptional immune rheostat. Predicted to successfully escape nonsense-mediated decay, and potentially driven by an activation-responsive NFAT promoter element, this isoform represents a highly plausible murine counterpart to human NKG2D-TR (4). While we rigorously frame this as a hypothesis requiring downstream experimental validation, the identification of this dynamic transcript adds an uncharacterized isoform to the murine NKG2D transcript landscape. These findings have important clinical and biological implications: they fill a critical cross-species gap in NKG2D biology, alter the interpretation of existing preclinical NKG2D-targeted immunotherapies, and highlight differentiation-coupled intron retention as a key mechanism that regulates the fine balance between graft-versus-tumor efficacy and graft-versus-host pathology.

### 4.1 Activation– and Differentiation-Associated Isoform Changes at the Klrk1 Locus

The central biological pattern revealed by our multi-dataset analysis is that *Klrk1-203* expression is not a stochastic artifact, but a tightly regulated transcriptomic switch intrinsically tied to T cell fate. We demonstrate that this isoform is fundamentally absent in healthy naive and early-effector CD8^+^ T cells (40), yet it robustly accumulates as cells mature into terminal effector and memory populations (41, 42). In ImmGen effector memory cells (LCMV day 180, n=1), *Klrk1-203* represents up to one-fifth of the total *Klrk1* pool (144.4 TPM), a substantial fraction similarly mirrored in GVHD CD4^+^ effector-memory cells (∼14%). Crucially, decomposing the retained-intron pool illuminates an orchestrated handoff in splicing logic during differentiation. In naive cells, the broader retained-intron fraction is dominated by *Klrk1-204* and *Klrk1-206*, with *Klrk1-203* undetectable. Therefore, the *total* retained-intron fraction is an inadequate marker of T cell state; the biologically significant signal is the specific, targeted induction of *Klrk1-203*. While we interpret the exact magnitude of this effect with appropriate bioinformatic caution, given that separating *Klrk1-203* from the co-retained *Klrk1-204* inherently relies on model-based quantification that short reads cannot independently resolve, the qualitative trajectory is unmistakable.

This nuanced remodeling is further supported by the GSE203167 post-transplant dataset. Here, total *Klrk1* is heavily induced and Klrk1-203 increases in absolute terms (∼5-fold from a near-zero baseline), although this isoform-specific change did not reach statistical significance. In GSE203167, the only dataset with sufficient replication for formal testing, a Dirichlet-multinomial differential transcript-usage test indicated significant overall Klrk1 isoform remodeling post-transplant (global p < 10⁻⁵; WT pre-vs-post-transplant p = 0.003). While *Klrk1-203* remains a minor proportional fraction (∼1.0%) in this specific acute setting, and thus did not reach statistical significance for specific usage change (p = 0.11–0.29), the overall retained-intron fraction significantly contracted (p = 0.02) driven by the selective collapse of *Klrk1-204*. This confirms that severe immunological stress triggers active, locus-wide isoform remodeling rather than a simple static scaling of expression.

Across 50 biological samples, the highly reproducible signal is the precise induction of *Klrk1-203* coinciding with effector and memory maturation. This dynamic is definitively resolved by the GSE119943 LCMV dataset: *Klrk1-203* remains entirely undetectable in highly proliferating, activated day-4.5 early effector cells, yet is robustly induced by day 7 in both SLEC and memory-precursor populations. Ultimately, this convergent observation points that *Klrk1-203* induction is not a mere byproduct of initial TCR engagement or general cellular stress, but a deliberate, developmentally regulated hallmark of terminal effector differentiation and memory commitment.

### 4.2 A Candidate NMD-Escaping, NKG2D-TR-Related Isoform

Cytotoxic responses mediated by NKG2D are extraordinarily potent and potentially tissue-destructive, calling for tight regulatory safeguards (43). We propose that the *Klrk1-203* retained-intron isoform functions as a vital post-transcriptional brake on murine NKG2D-mediated cytotoxicity. This hypothesis aligns conceptually (and, as our data now demonstrates, structurally) with the human NKG2D-TR splice variant, a dominant-negative isoform generated via activation-induced retention of intron 4 in the human *KLRK1* gene(18). While NKG2D-TR was originally identified as a human-specific regulatory mechanism, our structural mapping suggests that murine T cells employ a remarkably analogous strategy(18). The retained intronic sequence in *Klrk1-203* maps to a functionally identical location (the junction between the transmembrane and ligand-binding domains). If translated, this would produce a truncated product that retains the transmembrane segment through which adaptor association occurs, while lacking the annotated ligand-binding domain. In human NKG2D, assembly with DAP10 is mediated by basic arginine residues in the receptor transmembrane helix that form a three-helix interface with the acidic aspartate residues of the DAP10 dimer, and an isolated NKG2D transmembrane peptide is sufficient for this assembly(44). The predicted Klrk1-203 product retains the complete murine transmembrane helix (residues 67– 89), including both basic residues within it, and is therefore predicted to retain the structural capacity for DAP10 association. Sequence retention does not, however, establish membrane insertion, correct topology, stable expression, or actual adaptor binding, all of which require direct experimental testing. Topologically, this arrangement is comparable to human NKG2D-TR, which reduces functional NKG2D surface density both by competing for DAP10 and by promoting intracellular retention of the full-length receptor(18).

The profound significance of this finding lies in its regulatory logic. Although the specific intronic sequences retained differ between mice (intron 4 in *Klrk1-203*, interrupting the transmembrane and lectin-like domains) and humans (intron 4 of *KLRK1*, achieving identical topology), the overall strategy appears comparable: T cells actively couple the transcriptional upregulation of a dangerous activating receptor with simultaneous post-transcriptional dampening. While widespread intron retention is broadly recognized as a mechanism to tune transcriptomes and regulate hematopoietic processes like granulocyte differentiation, our findings extend this paradigm directly to the precise control of cytotoxic immune checkpoints. Given its robust induction to 144.4 TPM in CD8^+^ effector cells from a strict zero baseline in naive cells, *Klrk1-203* cannot be dismissed as transcriptional noise. Instead, we propose that this NMD-escaping transcript represents a potent, built-in rheostat. If even a fraction of this stable transcript pool is translated, it could compete for DAP10, functionally capping NKG2D signaling and preventing unchecked immunopathology.

For any alternative transcript to exert physiological influence, it must first survive cellular RNA surveillance. Two structural features are compatible with Klrk1-203 escaping degradation, though neither has been tested experimentally. First, the transcript is rigorously predicted to escape nonsense-mediated decay (NMD) under the canonical 50-nucleotide rule. Because its in-frame stop codon (positions 475 to 477) lies in the terminal exon (179 nucleotides downstream of the last exon-exon junction) it easily clears the ∼50-nucleotide threshold required to evade the NMD machinery. Second, the canonical *Klrk1-201* start codon is perfectly preserved at transcript position 112 within a robust Kozak context, and the longest unbiased open reading frame (ORF) initiates precisely at this site.

Together, this architecture dictates a stable transcript that is fully competent for translation using the exact same initiation machinery as full-length NKG2D(18). These structural realities leave three non-exclusive biological fates: nuclear retention to regulate local chromatin or transcription, cytoplasmic accumulation as a stable regulatory non-coding RNA, or direct translation into a truncated, dominant-negative protein. Given the striking structural overlap between the predicted *Klrk1-203* product and human NKG2D-TR, the translational hypothesis is uniquely compelling. We acknowledge that the algorithmic tool CPAT scores this transcript as non-coding (probability 0.158, below the 0.44 cutoff), a legitimate caveat that must be considered. However, because such predictive models are heavily calibrated for genome-wide differentiation of full-length mRNAs from long non-coding RNAs, they notoriously penalize legitimate short or truncated ORFs. Ultimately, the retention of the canonical initiation site and the topological similarity of human NKG2D-TR elevate the translation of *Klrk1-203* from a mere bioinformatic possibility to a testable hypothesis that requires ribosome-level resolution.

While our transcriptomic and structural analyses construct a compelling regulatory blueprint, definitively unmasking the functional fate of *Klrk1-203* requires targeted experimental execution. This computational framework provides the precise roadmap required for molecular validation: (Ribo-seq)(45), to confirm active engagement of the translation machinery at the preserved ORF, subcellular fractionation RNA-seq to ascertain whether the transcript is cytoplasmically exported or operates via nuclear retention, and high-resolution mass spectrometry to physically capture the predicted truncated NKG2D protein. Should translation be confirmed, rigorous functional assays evaluating DAP10/DAP12 sequestration in heterologous expression systems will be essential to conclusively demonstrate its dominant-negative capacity. Rather than serving as a generic catalog of future directions, our data provide the exact biological rationale and spatiotemporal coordinates required to execute these experiments successfully. By pinpointing precisely where, when, and how this transcript acts, this work sets the stage for the definitive experimental discovery of a native post-transcriptional brake on murine NKG2D.

The principal structural uncertainty in this study is the 3′ terminus of the annotated Klrk1-203 model, on which the NMD prediction depends. As detailed in the Discussion, the intron-4 retention event itself is supported directly by our alignment-level data, whereas the position at which these transcripts terminate rests on a single EST-derived transcript model. Targeted 3′-end mapping, by 3′ RACE from within exon 4 or by long-read sequencing of activated CD8⁺ T cell RNA, would resolve this and would simultaneously separate Klrk1-203 from the co-retained Klrk1-204; we regard it as the single most informative next experiment at this locus.

Retained-intron transcript models are frequently built from incomplete cDNA evidence. ENSMUST00000137660 carries transcript support level 3, indicating support from a single EST rather than from a full-length transcript, and the annotated model leaves only 24 nucleotides downstream of the predicted stop codon, which is short for a polyadenylated mRNA. We additionally found no canonical polyadenylation signal within 40 nucleotides of the annotated terminus. If the annotated terminus is instead an artifact of incomplete transcript support, the underlying molecule would represent a partially spliced intermediate that continues into the downstream exons; in that case the stop codon would no longer occupy a terminal exon and the transcript would be predicted to be NMD-sensitive rather than NMD-resistant. Distinguishing these possibilities requires 3′-end mapping, and we regard it as a prerequisite for any protein-level interpretation of this transcript. We also note the alternative that intron-4-retaining Klrk1 transcripts are nuclear-detained rather than exported, a widespread mode of post-transcriptional regulation that would be compatible with our expression data without invoking translation.

### 4.3 The TCF-7 Observation and Regulatory Inputs

The GSE203167 post-transplant dataset initially presented an apparent paradox: while absolute *Klrk1-203* expression increased during differentiation, the *overall* retained-intron percentage precipitously declined. Deconstructing this transcript pool resolved the contradiction. The decline was driven entirely by the selective collapse of *Klrk1-204* (the dominant pre-transplant background transcript) coupled with a massive induction of the canonical *Klrk1-201* isoform. *Klrk1-203* itself captured a rapidly expanding share of this shrinking retained-intron pool (surging from ∼4% to ∼29% in wild-type cells). This striking divergence confirms that intron retention at the *Klrk1* locus is not a monolithic, co-regulated block of transcriptional noise, but rather a suite of independently managed transcripts operating on distinct kinetic timescales. Crucially, this delicate splicing balance proved highly sensitive to the broader transcriptional landscape. In *TCF-7* conditional knockout cells, canonical *Klrk1-201* hyper-dominates the transcript pool (constituting roughly 63% of total *Klrk1* post-transplant). TCF-7 is the master regulator of T cell stemness, widely recognized for actively repressing terminal effector differentiation and limiting exhaustive cytotoxicity. The pronounced shift toward the canonical, highly cytotoxic *Klrk1* isoform in the absence of TCF-7 raises a provocative hypothesis: TCF-7 may preserve less cytotoxic, stem-like CD8+ T cell states partially by enforcing a post-transcriptional splicing balance that favors inhibitory or retained-intron variants over full-length activating receptors.

Furthermore, the transition from a *Klrk1-204/206*-dominated naive background to an actively induced *Klrk1-203* state in differentiated cells indicates that transcript accumulation at this locus reflects active splice-site selection rather than mere transcriptional leakage. While global intron retention and CD28-driven alternative splicing have been broadly documented as mechanisms shaping T cell activation, our data anchor this regulatory paradigm to a specific, critical immune checkpoint. Ultimately, these findings support Klrk1 intron retention as a differentiation-associated event for post-transcriptional diversification, one that appears intimately wired into the core transcriptional networks dictating T cell fate.

### 4.4 A Two-Step Model for *Klrk1* Regulation

To fully map the regulatory architecture of the *Klrk1* locus, we must uncouple initial transcription from post-transcriptional processing. Our *in silico* promoter analysis identifies a high-confidence, shared NFAT binding motif located precisely 134 bp upstream of the canonical *Klrk1* transcription start site (33). This places TCR-driven calcineurin-NFAT signaling (a pathway that is intrinsically linked to NK-receptor expression on CD8^+^ T cells) as the likely primary trigger for locus-wide transcription (46). The relevant point here is how this transcriptional input integrates with our splicing data. An NFAT-driven transcriptional burst would non-specifically generate pre-mRNA for the entire locus, driving both canonical and retained-intron precursor transcripts at the same time. Therefore, the highly specific, temporally restricted emergence of *Klrk1-203* cannot be explained by promoter activation alone; it demands a distinct, second-tier post-transcriptional filter. While this two-step model (TCR-driven bulk transcription followed by differentiation-gated alternative splicing) requires formal experimental validation, it elegantly explains a critical biological paradox: how T cells can rapidly and broadly mobilize an activating receptor upon antigen encounter, while retaining the precise ability to independently tune and throttle its output as they mature into memory states.

### 4.5 Technical Controls

Computational transcriptomics of highly homologous isoforms is notoriously susceptible to artifacts, demanding rigorous technical validation. Anticipating these challenges, we applied several orthogonal controls. The technical controls described in the Results address several distinct artifact classes: the retained-intron signal was also not higher in the one rRNA-depleted dataset than in poly(A)-selected datasets, arguing against library chemistry as its source. The low retained-intron signal at highly expressed housekeeping genes (*Actb*, *Gapdh*), the direct junction-spanning PIR validation extracted from raw alignments, and the Sashimi visual confirmations collectively argue against genomic DNA contamination or incomplete library size selection as the source of the signal. Furthermore, we demonstrated that stringent sequence– and GC-bias correction is mandatory for accurate *Klrk1* quantification; omitting it in preliminary tests artificially inflated minor overlapping transcripts (e.g., *Klrk1-202*) from 0 to 52 TPM. Applying this control pipeline across all 50 samples supports a gene-specific biological signal rather than background noise. Together, these controls make a purely technical origin for the Klrk1 retained-intron signal unlikely, although they cannot exclude residual quantification bias at this locus. Klrk1-203 usage was also not predicted by total Klrk1 abundance within the LCMV dataset, arguing against an expression-dependent quantification artifact.

### 4.6 Translational Implications for Immunotherapy

Understanding species-specific splicing architectures is not merely an academic exercise; it is the critical linchpin for the translational pipeline of NKG2D-targeted therapies. Murine models remain the primary preclinical platform for evaluating NKG2D blockade and CAR-T cell strategies in both GVHD and cancer immunotherapy. The murine NKG2D system is already functionally complex due to the existence of the alternative-promoter NKG2D-L and NKG2D-S isoforms, which dictate differential DAP10/DAP12 coupling. If *Klrk1-203* operates as the dominant-negative regulator our structural analysis predicts, it introduces a profound variable: murine models of NKG2D-targeted GVHD and tumor therapy already harbor a dynamic, endogenous negative feedback loop that fundamentally mirrors the human NKG2D-TR system. This striking cross-species convergence may warrant consideration in the interpretation of NKG2D biology. Particularly in the context of antibody-based blockade or genetic NKG2D ablation strategies, this endogenous splicing rheostat could actively complement or severely confound therapeutic interventions. Extrapolating murine efficacy to human trials without consideration of this intrinsic regulatory brake risks a fundamental misinterpretation of NKG2D biology.

### 4.7 Methodological Constraints and Reproducibility

We acknowledge several methodological constraints, which our multi-layered validation strategy was explicitly designed to mitigate. First, per-condition sample sizes across these datasets are small (e.g., *n* = 1 Tn and *n* = 3 Tem in the GVHD cohort), and the GSE203167 dataset exhibited an underpowered, non-significant *Klrk1-203*-specific signal despite massive global isoform remodeling. Second, read length presents a fundamental bioinformatic constraint. Separating highly homologous transcripts inherently relies on model-based quantification (Salmon) to resolve overlapping reads. We addressed this by overlaying an isoform-inference-free, read-level assessment of the exon 4–intron 4 boundary directly from the raw alignments. While this alignment-level signal reports intron-4 retention as a whole, and cannot independently separate *Klrk1-203* from the co-retained *Klrk1-204*, its tight convergence with our transcript-level estimates is difficult to reconcile with short-read ambiguity as the source of our signal.

Furthermore, our study is strictly transcriptomic and *in silico*. Direct experimental measurement is absolutely required to confirm transcript stability, translation of the predicted open reading frame, functional DAP10 sequestration, and true physiological NFAT occupancy at the *Klrk1* promoter. The datasets analyzed here derive from a small number of related backgrounds (C57BL/6, and B10.D2 into BALB/c for the GVHD model), leaving broader strain and age effects to be mapped. However, the ultimate strength of this study derives from independent reproducibility rather than isolated within-dataset significance testing. A consistent Klrk1-203 induction pattern emerges across four datasets generated by separate laboratories on different sequencing platforms, spanning viral infection, chronic GVHD, and allogeneic transplantation. We note that the ImmGen and GSE119943 effector and memory populations both derive from LCMV infection and are therefore not fully independent of one another. A biological pattern that resiliently holds across entirely unrelated experimental contexts is profoundly harder to attribute to chance, or to a single technical batch effect, than a highly significant *p*-value derived from a single isolated cohort. Ultimately, these overlapping transcriptomic and structural confirmations elevate *Klrk1-203* from an obscure bioinformatic annotation to a high-priority biological target.

The principal structural uncertainty in this study is the 3′ terminus of the annotated Klrk1-203 model, on which the NMD prediction depends.

Transcript identifiers and versions are as annotated in Ensembl release 115, the release used to build the quantification index. The Klrk1 annotation has been updated in subsequent releases, including the addition of a further protein-coding transcript model (Klrk1-207, ENSMUST00000573601) that was not present in release 115 and is therefore not included in the reporting units defined here.

### 4.8 Future Directions and Concluding Remarks

Two further controls were beyond the scope of the present analysis and are acknowledged as limitations. First, we did not perform simulation-based validation in which reads are generated from a defined ground-truth mixture lacking Klrk1-203 and passed through the same pipeline; such an experiment would formally bound the rate at which model-based apportionment can assign signal to Klrk1-203 in its absence, particularly at high total Klrk1 expression. Second, all quantification used transcriptome-only Salmon indices. Although these showed high concordance with a decoy-aware index in a test sample, decoy-aware indices are specifically designed to limit the misassignment of intronic and genomic reads, which is the artifact class most relevant to retained-intron quantification; a systematic decoy-aware reanalysis remains to be performed. Both analyses are straightforward extensions of the present pipeline, and we regard them as the natural next computational step.

The identification of Klrk1-203 as a dynamic, differentiation-coupled transcript establishes a definitive roadmap for translating murine NKG2D biology into precision immunotherapies. First, extending this isoform-level resolution to statistically powered in vivo tumor-immunity models will be critical to confirm these kinetic dynamics in a therapeutic context. Applying advanced differential isoform frameworks (such as IsoformSwitchAnalyzeR) to robust datasets (particularly targeted tumor-immunity models) will establish whether the magnitude of Klrk1-203 induction directly correlates with anti-tumor efficacy or host cytotoxicity. Furthermore, large-scale cross-species transcriptomic comparisons must be deployed to determine the precise extent to which this intron-retention program is shared between murine Klrk1 and human KLRK1.

Functionally, resolving the precise cellular fate of *Klrk1-203* now represents a critical frontier in immune checkpoint biology. Our computational architecture predicts a stable, translationally competent transcript that yields a 121-amino-acid truncated product, one that retains its cytoplasmic and transmembrane domains but entirely lacks the ligand-binding ectodomain. The immediate experimental mandates are clear: ribosome profiling (Ribo-seq) of activated CD8+ T cells to confirm active ribosomal engagement at the predicted *Klrk1-203* initiation site, followed by mass spectrometry of immunoprecipitated NKG2D species to physically validate protein-level output. If translated, heterologous expression assays must be utilized to confirm DAP10 co-immunoprecipitation and the subsequent reduction of functional surface NKG2D. Critically, targeted *in vivo* depletion or stabilization of *Klrk1-203* using antisense oligonucleotides (ASOs) or CRISPR-based editing would allow us to therapeutically titrate NKG2D signaling, providing a novel approach to maximize graft-versus-tumor (GVT) efficacy while aggressively suppressing GVHD pathology.

Mechanistically, the upstream regulation of this splicing event remains to be defined. Combining these transcriptomic profiles with chromatin accessibility assays (ATAC-seq, CUT&Tag) will provide essential insights into how TCR-driven epigenetic remodeling coordinates with splice-site selection at the *Klrk1* locus. Our observation in *TCF-7* conditional knockout cells (where canonical isoform dominance drastically increases at the expense of the retained-intron fraction) generates a provocative, testable hypothesis: TCF-7 actively maintains a protective splicing balance, and its loss unleashes productive, potentially exhaustive cytotoxic output. Concurrently, rigorous functional interrogation of the predicted proximal NFAT element via luciferase reporter assays, site-directed mutagenesis, and chromatin immunoprecipitation will determine if TCR-driven calcineurin signaling serves as the master transcriptional ignition switch for this entire locus. By mapping the structural and regulatory parallels between murine *Klrk1-203* and human NKG2D-TR, this study positions orchestrated intron retention not merely as a biological curiosity, but as a central, targetable axis of immune regulation. This post-transcriptional checkpoint, once unlocked, fundamentally reshapes our understanding of murine preclinical models and provides a powerful mechanistic blueprint for uncoupling protective immunity from destructive immunopathology in next-generation immunotherapies.

## Data and code availability statement

All primary data are publicly available from NCBI GEO (GSE147371, GSE109125, GSE203167, and GSE119943). Salmon quantification files, R analysis scripts, HISAT2 alignment commands, CPAT inputs, and all intermediate analysis logs will be deposited on request.

## Author contributions

IHT: conceptualization, data curation, data analysis, investigation, methodology, software, visualization, writing – original draft. MK: conceptualization, data analysis, methodology, funding acquisition, project administration, resources, supervision, writing – review and editing. All authors contributed to the article and approved the submitted version.

## Acknowledgements

The authors thank all members of the Karimi Laboratory for their helpful discussions and technical support throughout this study.

## Ethics statement

This study analyzed publicly available, de-identified RNA-seq datasets and required no new human or animal ethics approval. The animal work underlying GSE203167 was performed under IACUC protocol number 433, approved by the Institutional Animal Care and Use Committee of SUNY Upstate Medical University.

